# Including fitness and health proxies can alter our understanding of habitat selection

**DOI:** 10.1101/2025.11.13.688337

**Authors:** Marie Auger-Méthé, Fanny Dupont, Alyssa Eby, Kyle H. Elliott, Nigel E. Hussey, Devin A. Lyons, Marianne Marcoux, Allison Patterson, Shabnam Shadloo, Courtney R. Shuert

## Abstract

Habitat selection analyses, which discern the environmental conditions individuals select, often inform conservation planning. Through a literature review, we demonstrate that recent habitat selection studies rarely include fitness and health information. With a simulation study, we show that ignoring such information could support the protection of sink habitats. Our case studies demonstrate how health and fitness proxies can modify our understanding of habitat selection: (1) incorporating mass gain of thick-billed murres shows the energetic benefit of areas deemed secondary by a naive resource selection function; (2) including number of chicks in a step selection function (SSF) exposes the complex relationships glaucous-winged gulls have with landscapes impacted by humans; and (3) including external signs of trauma in the movement kernel of SSFs demonstrate others ways in which narwhal distribution can be altered. We urge movement ecologists to collect and use health and fitness data to improve ecological inference and conservation action.

## 1 Introduction

Habitat selection has been a central topic of research for decades due to its importance in understanding the environmental conditions that explain the distribution of animals (Northrup et al., 2022). Its analysis has long been motivated by the theoretical argument that habitat-use patterns result from selection pressures on the survival and reproduction of animals (Boyce and McDonald, 1999). This link to natural selection processes allows ecologists to infer that habitats selected with higher intensity are perceived as having higher fitness payoffs (Boyce and McDonald, 1999; Matthiopoulos et al., 2023; Northrup et al., 2022), and thereby allows managers to focus on conserving habitats that are strongly selected by animals. Given the increasing use of GPS tags and other tracking devices (Nathan et al., 2022), resource selection functions (RSFs) and step selection functions (SSFs) have become important and increasingly used tools to quantify habitat selection and inform management plans (Florko et al., 2025; Lemieux Lefebvre et al., 2018; Northrup et al., 2022). However, both density and intensity of use can be misleading indicators of habitat quality; thus, habitat selection analyses can lead to flawed conservation actions (Aldridge and Boyce, 2008; Van Horne, 1983).

Fitness increases with habitat selection under a set of simple, but commonly violated, assumptions (Matthiopoulos et al., 2023). Habitat selection models hinge on the expectation that animals distribute themselves with an ideal free distribution (Fretwell and Lucas, 1969), have perfect knowledge of the environment, and can respond to it immediately (Gaillard et al., 2010; Matthiopoulos et al., 2023). However, social interactions can prevent subdominant individuals from entering high-quality habitats and relegate them to sink habitats where they have a low probability of surviving and reproducing (Van Horne, 1983). In addition, environmental conditions are often stochastic, and an animal’s knowledge and perception of its surroundings are inherently partial. Individuals may use cues of habitat quality that are no longer reliable due to environmental changes, with ecological traps caused by rapid human-induced changes potentially creating large mismatches between habitat selection and fitness (Robertson et al., 2013; Schlaepfer et al., 2002). Similarly, site fidelity can result in animals using habitats that are no longer advantageous, a maladaptive behaviour that is likely to increase with rapid human-induced changes (Merkle et al., 2022; Van Horne, 1983). Many other factors may explain mismatches between habitat selection and fitness, including multi-generational delays in colonisation processes (Matthiopoulos et al., 2023).

Many have argued that quantifying habitat quality in terms of its contribution to fitness cannot rely solely on habitat selection and must include data on population dynamics or individual performance (DeCesare et al., 2014; Gaillard et al., 2010; Lemieux Lefebvre et al., 2018; Matthiopoulos et al., 2015; Van Horne, 1983). For example, Matthiopoulos et al. (2015) proposed a framework that links habitat selection to population growth, thus disentangling apparent habitat suitability from the true contribution of habitat to fitness. Although this approach is powerful, it requires multiple population time series, each linked to animal space-use data. Thus, when historical time series are unavailable, it may be challenging to implement this framework fast enough to solve many urgent conservation problems.

It has long been recognised that habitat quality can be quantified by combining space use with basic vital rates such as survival and offspring production (Van Horne, 1983), with important papers (e.g., DeCesare et al., 2014; Gaillard et al., 2010) re-emphasising the need for habitat selection analysis to be linked to fitness-related data. A fruitful research area has been the combination of habitat selection and survival analyses, resulting in the development of methods to quantify instantaneous mortality risks and long-term drivers of survival (e.g., DeCesare et al., 2014; Eisaguirre et al., 2025; Moeller et al., 2025; Poulton et al., 2024). However, adult survival data can be challenging to gather, especially for long-lived species, and other components of individual fitness or proxies of individual performance can be measured (e.g., annual reproductive success, energy gain; Gaillard et al., 2010).

Although animals show high levels of individual variation in habitat selection, and it is recognised that ignoring this variation can mask important relationships (Bastille-Rousseau and Wittemyer, 2019; Leclerc et al., 2016), most analyses that include this variation either only account for it statistically (e.g., Muff et al., 2020) or attempt to explain it with non-fitness related characteristics (e.g., sex; Bastille-Rousseau and Wittemyer, 2019; Winter et al., 2024). A recent review on habitat selection (Northrup et al., 2022) emphasised that we could learn more by treating individual variation as a feature to be explored, rather than a nuisance. Here, we further argue that explaining individual variation in habitat selection using proxies of individual fitness or health can help us gain crucial insights into the causes and consequences of this variation and, when appropriate measures of individual performance are chosen, get us closer to quantifying habitat quality.

Using a literature review, we assess whether recent habitat selection studies include fitness and health information. We then use simulations to demonstrate the potential biases that can arise when these proxies are ignored and show the reliability of a simple, but underused, method to integrate such proxies in RSFs and SSFs. Finally, we use three animal movement case studies (Figure 1) to demonstrate how a variety of fitness and health proxies can be used to provide habitat selection insights that would otherwise be hidden.

**Figure 1:**
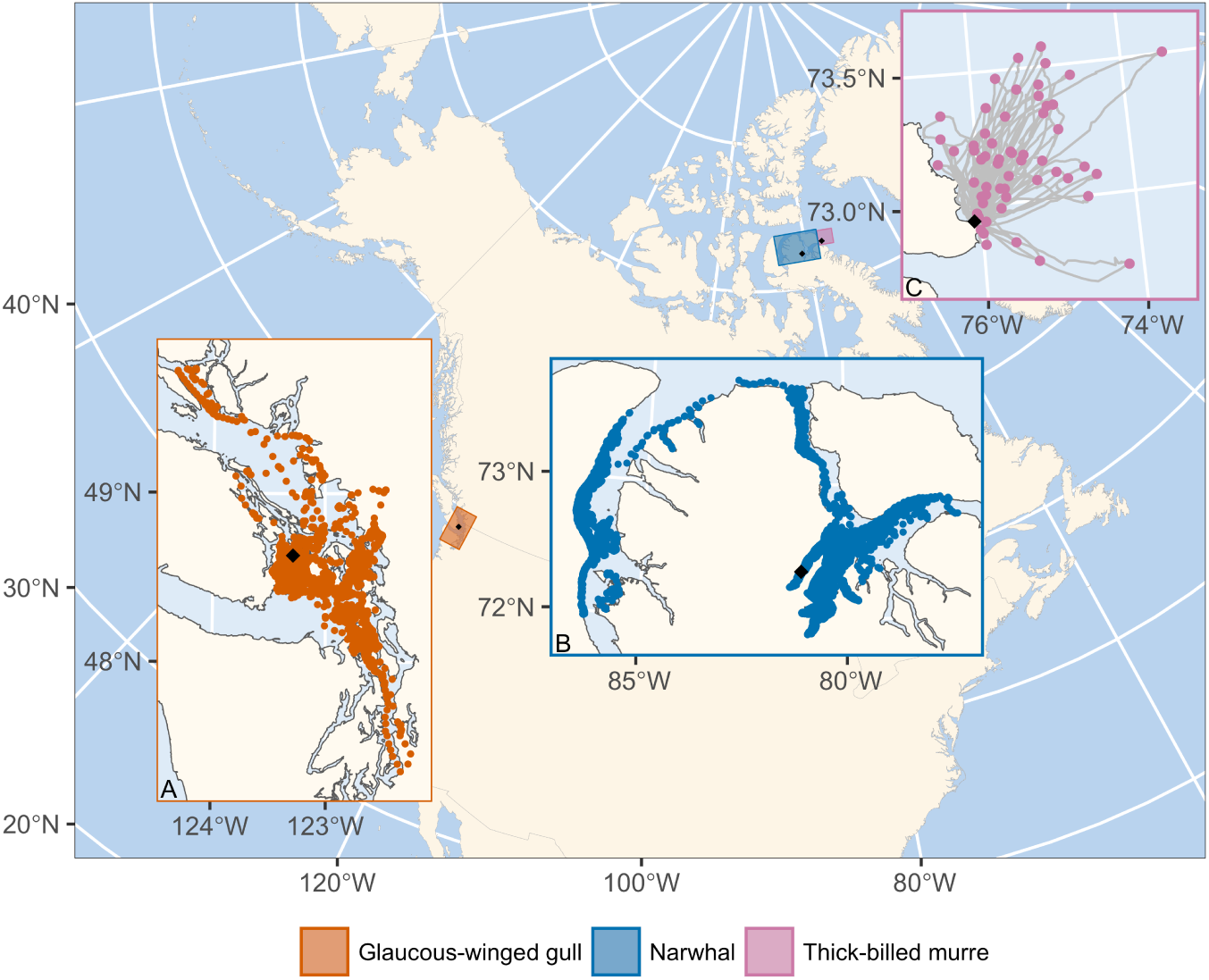
Field site location of the three case studies (black diamonds) and the location data used for each analysis (coloured points). Panel A shows daytime GPS locations of glaucous-winged gulls (orange points). Panel B shows pre-processed Fastloc GPS data of narwhals (blue points). Panel C shows foraging trips of the thick-billed murres (grey lines). For each trip, the purple point identifies the location furthest from the colony.

## 2 Methods

### 2.1 Literature review

We conducted a literature review to assess the frequency with which recent habitat selection studies include fitness or health information. We queried Web of Science using the search terms: (resource selection function OR step*selection function OR habitat selection analys*) AND (telemetry OR tag* OR collar* OR GPS OR Global Positioning System).

We searched for English peer-reviewed articles published between April 29, 2024 and April 28, 2025 that performed a habitat selection analysis based on a used-available approach (*sensu* Matthiopoulos et al., 2023; e.g., RSF, SSF, inhomogeneous Poisson point process) to GPS tracking data of non-human animals. We were interested in whether individual characteristics were accounted for; thus, we only kept papers that included more than two individuals in their habitat selection analysis and that contained sufficient information to assess how individual characteristics were included. We extracted information that allowed us to compare the use of fitness and health proxies to that of other individual characteristics (e.g., sex) and assess how often habitat selection analyses are used to make conservation or management recommendations. See Supporting Information for further details.

### 2.2 Simulations

We used simulations to demonstrate potential biases that could occur in habitat selection analyses when ignoring individual differences in fitness or health, and to verify that the simple method we propose can accurately estimate parameters. For each run, we simulated 30 individuals, each with one of three health proxy values: -1, 0, and 1, representing inferior, average, and superior health, respectively. To explore whether the proportion of inferior, average, and superior health individuals in the sample affects the estimated selection parameters, we created simulations for three different inferior:average:superior ratios: an inferior-skewed sample ‘15:10:5’, a balanced sample ‘10:10:10’, and a superior-skewed sample ‘5:10:15’.

For each simulation run, we created a 100 x 100 raster representing an environmental covariate by sampling a Gaussian random field with scale 0.9 and standard deviation 0.1 using the R package prioritizer (Hanson et al., 2023). For each individual, we randomly sampled 50 cells of this raster, with the probability of selecting cell *i* defined as:

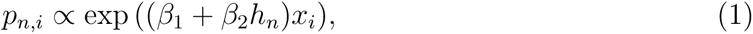

where *x_i_* is the value of the environmental covariate for cell *i*, *h_n_* is the value of the health proxy for the *n^th^* individual, *β*_1_ represents the selection coefficient for the covariate when the health proxy is 0 and *β*_2_ represents an interaction between the selection of the environmental covariate and the health proxy.

We explored two scenarios that could result in problematic conclusions. Scenario A (*β*_1_ = 0.5, *β*_2_ = 1) represents a situation in which individuals in superior health select strongly for high values of the covariate (*β*_1_ + *β*_2_*h_n_* = 1.5), individuals in average health select moderately for high values of the covariate (*β*_1_ + *β*_2_*h_n_* = 0.5), and individuals in inferior health avoid areas with high values of the covariate (*β*_1_ + *β*_2_*h_n_* = −0.5). This scenario could represent a situation where individuals in superior health restrict access to a highly beneficial resource. Scenario B (*β*_1_ = 0, *β*_2_ = −1) represents a situation where individuals in superior health avoid high values of the covariate (*β*_1_ + *β*_2_*h_n_* = −1), individuals in average health are indifferent to this covariate (*β*_1_ + *β*_2_*h_n_* = 0), and individuals in inferior health select for this covariate (*β*_1_ + *β*_2_*h_n_* = 1). This scenario could represent a situation where the environmental covariate is associated with a disease, leading those selecting for it to become sick. We simulated each combination of scenario and proxy ratio 100 times for a total of 600 simulations.

For each simulation, we estimated the parameters of two RSFs, which are commonly used functions that quantify the relative strength with which a location is selected based on environmental covariates (see Supporting Information, Florko et al., 2025; Northrup et al., 2022). The baseline RSF included only the relationship with the environmental covariate, while the proxy-informed RSF represented the selection process used to simulate the data and, thus, also included the relationship between the environmental covariate and the health proxy. Our estimation procedure used the common use-available design where observed animal locations (i.e., *used locations*) are compared to locations sampled within the area assumed to be available to the animal (i.e., *available locations;* Florko et al., 2025; Northrup et al., 2022). Specifically, we used the method described by Muff et al. (2020), which approximates the model likelihood by sampling, for each used location of each individual (*y_n,j_*= 1 where *j* = 0), *J* available locations (*y_n,j_*= 0 for *j* = 1*, . . . , J* ) and fitting a generalised linear mixed model (GLMM) via glmmTMB (Brooks et al., 2017). This method allows the inclusion of random effects, which can account for unexplained individual variation and the nested nature of many datasets. Specifically, the GLMM used to estimate the parameters of the baseline RSF modelled the probability that a location *y_n,j_* with covariate value *x_n,j_*is used, *π_n,j_*, as:

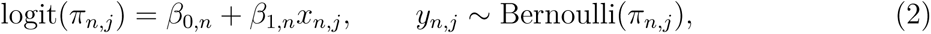

where logit represents the logistic link function, the intercept is a random effect with a fixed standard deviation (*β*_0_*_,n_* ∼ N(*β*_0_, 10000^2^)), and the selection coefficient is a random effect for which we estimate the standard deviation (*β*_1_*_,n_* ∼ N(*β*_1_*, σ*^2^*_β_*_1_); Muff et al., 2020). To estimate the parameters of the proxy-informed RSF, we simply added an interaction with the health proxy:

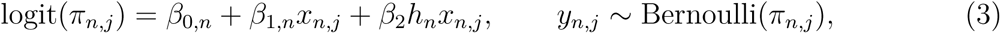

where *β*_2_ represents the relationship between selection and the health proxy, and is not modelled as a random effect because *β*_1_*_,n_* already accounts for unexplained individual variation in selection. We did not include a term with the health proxy by itself (e.g., *β*_3_*h_n_*), as its coefficient would simply quantify sample size differences across groups (Northrup et al., 2022). For both RSFs, we sampled 100 available locations per used location using the R package amt (Signer et al., 2019) and used a weighted logistic regression (weight of available = 1000 and used = 1; Muff et al., 2020). We compared the two RSFs using the Akaike Information Criterion with small-sample correction (AICc, Burnham and Anderson, 2002).

We also conducted SSF simulations. Step selection functions differ from RSFs in that they account for the movement of the animal and limit the availability to areas that can be reached during two consecutive locations (see Supporting Information, Florko et al., 2025; Thurfjell et al., 2014). The simulated movement model was based on the integrated SSF framework of Avgar et al. (2016), which models the movement as a combination of a habitat selection function and a selection-free movement kernel that describes the movement of the animal in the absence of habitat selection. We used the same selection coefficients as scenarios A and B, and the same three health proxy ratios. We estimated the parameters of a baseline SSF that included only the environmental covariate and a proxy-informed SSF that included an interaction between the covariate and the health proxy using the method of Muff et al. (2020). See Supporting Information for more details.

To verify that AICc would correctly select the baseline model when there are no relationships with the health proxy, we created additional scenarios where the baseline RSF and SSF were simulated.

### 2.3 Case studies

We used three simple case studies to demonstrate the ease with which various health and fitness proxies can be included in habitat selection analyses. In all case studies, we compared habitat selection models that included a proxy to a baseline model that ignored it. As with the simulations (Section 2.2), we estimated the parameters of the models using the method of Muff et al. (2020). We compared the models using their difference in AICc values, ΔAICc, while considering the difference in the number of parameters estimated, Δ*k* (Sutherland et al., 2023). We also examined the coefficients associated with the proxy.

#### 2.3.1 Case study 1: including proxy of mass gained

Case study 1 used movement data from 54 thick-billed murres (*Uria lomvia*; hereafter murres) to demonstrate how habitat selection can be informed by a measure of energy gain: the rate of mass gain. Our dataset consisted of incubating adults tagged in July and August 2022 and 2023 at their nest in Nunavut, Canada (72°56.5500 N, -76°5.6880 W, Figure 1). As the murres were recaptured *<* 26 hrs later, the movement track of each individual, which was collected with a Technosmart Axy-trek mini tag programmed to collect GPS locations at 1 or 3 min intervals, represented a single foraging bout. To identify where they forage before returning, we chose the location farthest from the colony (Figure 1C). The murres were weighed to the nearest gram at capture and recapture. To account for the deployment duration on mass gained, we calculated the rate of mass gained (g/hr). Research was conducted under the McGill University Animal Care permit 2015-7599, Environment and Climate Change Canada (ECCC) scientific permit to capture and band migratory birds 10892, and Nunavut government permit 2022-024.

The baseline murre RSF model included only distance to colony as a covariate for selection, while the proxy-informed RSF also included an interaction with rate of mass gain, allowing this proxy to modify habitat selection. As there was one foraging location per individual, we did not include a random effect on distance to colony. We sampled 1000 available locations per used location from a study area that excluded areas on land and had a 125 km radius around the colony (maximum observed distance: 106 km). See Supporting Information for more details.

#### 2.3.2 Case study 2: habitat selection relationship with proxies of annual reproductive success

Case study 2 used the movement of 12 glaucous-winged gulls (*Larus glaucescens*; hereafter gulls) tagged on a relatively pristine island in British Columbia, Canada (48°38.0400 N, -123°17.2200 W, Figure 1) to demonstrate how habitat selection can be linked to proxies of reproductive success. The gulls were captured at their nest during the early incubation period (June 11-13). They were tagged with OrniTrack-E20 4G tags (Ornitela) that collected a GPS location every 15 min. To monitor the reproductive success of each individual, we banded them with a unique combination of Canadian Wildlife Service metal and coloured plastic leg bands and identified their nests with numbered flags. We surveyed nests every 6 ± 4 days and used the number of chicks (ranging from 0 to 3) on July 31 as our proxy of annual reproductive success. To examine foraging habitat during the period linked to this proxy, we used movement data from 1 day after capture until July 31. Gulls primarily forage during the day (Hayes and Hayward, 2020; Henson et al., 2004), thus we kept only daytime locations that were outside a 200 m buffer around the colony. Research was conducted under the University of British Columbia Animal Care permit A22-0045, ECCC scientific permit to capture and band migratory birds 10667Z.

The two SSFs explored included as covariates: (1) distance to shore, as this coastal species does not travel far inland or offshore; (2) distance to colony, as they are central-place foragers when nesting; (3) a binary ‘land’ variable representing whether they use terrestrial or marine environments (land = 1, ocean = 0); and (4) a human impact layer (Mu et al., 2021; Mu et al., 2022). The human impact layer is only for the terrestrial environment; thus, we included it as an interaction with the binary land variable. We added random effects on the selection coefficients associated with distance to shore, distance to colony, land, and the interaction between land and human impact. While the baseline model only included these covariates, the proxy-informed model included the number of chicks as an interaction with both distance to colony and the land by human impact interaction. We sampled 100 control locations per used location. See Supporting Information for more details.

#### 2.3.3 Case study 3: habitat selection relationships with old injuries

Case study 3 used the movement data of 14 narwhals (*Monodon monoceros*) to demonstrate how external signs of trauma can inform habitat selection. The narwhals were captured during the summer in Nunavut, Canada (72°21.1389 N, -81°05.855 W, Figure 1) and outfitted with tags (TDR10, Wildlife Computers, Inc.) that provided Fastloc GPS data. To account for the short-term effects of handling (Shuert et al., 2021), the migratory behaviour of narwhals (Shuert et al., 2022), and the irregular nature of Fastloc GPS for marine animals (Dujon et al., 2014), we pre-processed the movement data as in Shuert et al. (2025). This procedure resulted in summer movement tracks with a location every 1 hr. Before tagging, we performed a visual assessment of the physical condition of individuals and recorded the presence of healed scars. These scars indicate previous trauma, such as negative interactions with humans (e.g., gunshot wounds), other narwhals (e.g., tusk scars), and predators (e.g., tooth rakes; Auger-Méthé et al., 2010). All capture and tagging protocols were approved by the Animal Care Committees of Fisheries and Oceans and the University of British Columbia, and a License for Scientific Purposes was granted (permit #FWI-ACC-2016–030, AUP 40, FWI-ACC-2018–22, ACC A18-0179).

To assess whether healed scars should be included in the habitat selection function or the selection-free movement kernel, we explored four SSFs. In all models, distance to shore was the only environmental covariate. However, it was included in the selection function as the distance from shore to the locations at the end of steps, to represent narwhals choosing where to go, and in the movement kernel as the distance to the locations at the start of steps, to represent narwhals changing their speed based on how close they are to land. To incorporate distance to shore in the movement kernel, we included it as an interaction with step length and the natural logarithm of step length. We also added an interaction between the squared distance to shore and the natural logarithm of step length because the relationship was not linear. We added a random effect on the selection coefficient associated with distance to shore. The first SSF explored was the baseline model, which ignored healed scar information. The second SSF included the presence of healed scars only in the selection function as an interaction with distance to shore. The third SSF included the presence of healed scars in the movement kernel as a three-way interaction with step length and distance to shore. The fourth SSF combined the last two models and included the presence of healed scars in both the selection function and movement kernel. We sampled 100 control locations per used location. See Supporting Information for more details.

## 3 Results

### 3.1 Literature review

Of the 72 articles returned by our literature search, 82% (59/72) made statements about conservation and management based on the results of their analyses (Figure 2A). Of the 13 papers not making such statements, five focused on methodological developments. Although these results suggest that many ecologists use habitat selection analyses to inform conservation and management, only 4% (3/72) of the articles included a characteristic of individual health or fitness in their habitat selection analyses (Figure 2B), with only a few others (7/72) considering reproduction or survival via a focus on calving habitat (i.e., analyses restricted to pregnant females or those with neonates), kill sites, or by including a separate survival analysis (Figure 2B). Other individual characteristics were used more frequently (28%, 20/72), with sex being the most common (15/20).

**Figure 2:**
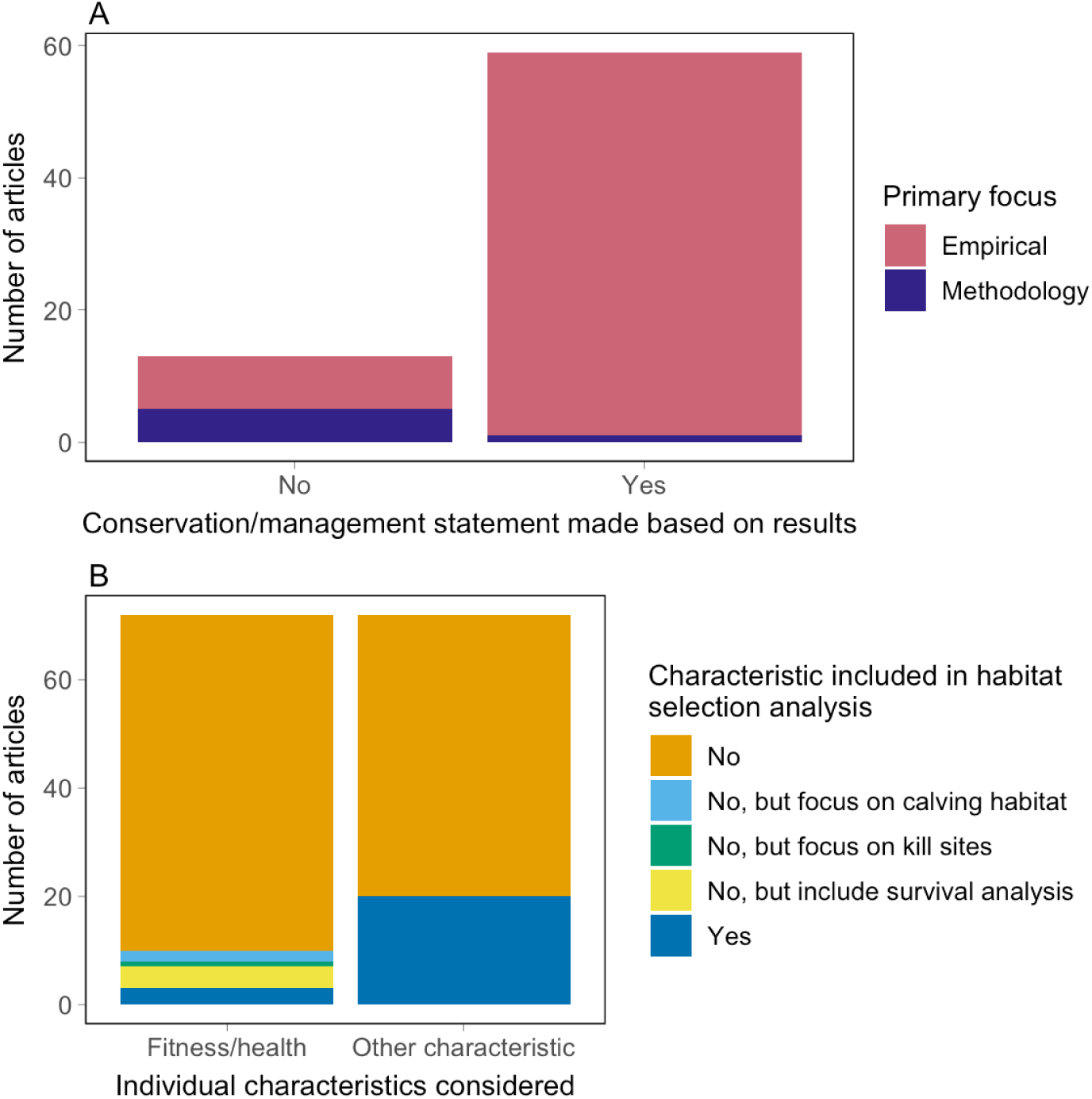
Bar plots representing the results from the literature review on habitat selection analysis. Panel A shows the number of recent articles that made a statement on conservation or management based on the results. Colours represent whether the articles were focused primarily on methodology (dark purple) or the empirical analysis (magenta). Panel B displays the number of recent articles that did (dark blue) or did not (orange) include individual characteristics in the habitat selection analysis, as well as articles that did not explore how fitness or health characteristics affect habitat selection, but that included related characteristics as either an additional survival analysis, detection of kill sites, or a focus on calving habitat.

### 3.2 Simulations

The simulations showed that when there is an imbalance in the number of healthy versus unhealthy individuals in the sample, applying a simple RSF or SSF can return biased results (Figures 3 and S1). For example, in scenario A, the estimates of the selection coefficient of the baseline RSF (*β*_1_ in equations 2) were negatively biased when there were more individuals in inferior health. In addition, their 95% confidence intervals overlapped with 0 in 100% of the simulations, which, under a null hypothesis testing approach, would be interpreted as evidence for a lack of selection of this covariate, even though individuals in superior health select for it strongly. In scenario B, the estimates of the selection coefficient of the baseline RSF (*β*_1_ in equations 2) were positively biased when there were more unhealthy individuals in the sample. In addition, their 95% confidence intervals excluded 0 in 97% of the simulations, which would be interpreted as evidence for selection for the environmental covariate, even though it is avoided by individuals in superior health.

**Figure 3:**
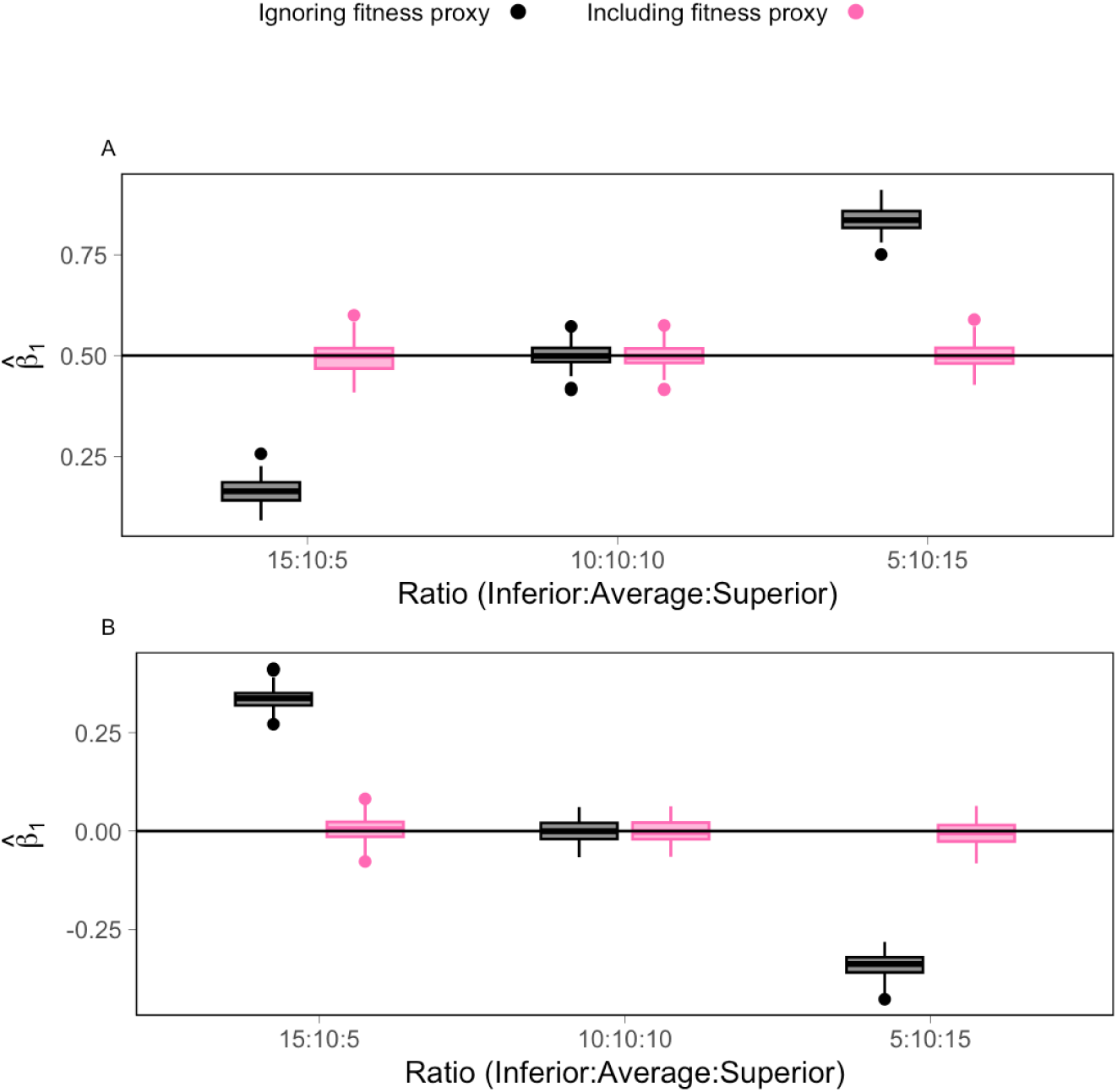
Boxplots of the estimated selection parameter *β*_1_ of an RSF under two scenarios and three ratios of individuals. The black horizontal lines represent the value used to simulate the data: scenario A, *β*_1_ = 0.5; scenario B, *β*_1_ = 0. Scenarios A and B represented, respectively, an increasing (*β*_2_ = 1) and decreasing (*β*_2_ = −1) selection for the covariate with health/fitness proxy condition. The black boxplots represent the estimates when we used an RSF that ignored fitness proxies, while the pink boxplots represent the estimates when the fitness proxy was included in the RSF as an interaction with the environmental covariate.

As detailed in the Supporting Information, when the simulations included a relationship with a health proxy, AICc always selected the proxy-informed models over baseline models, and these proxy-informed models returned unbiased parameter estimates. When we simulated baseline RSFs and SSFs that did not include a relationship with the health proxy, AICc wrongly selected the model that included the health proxy 9-12% of the time. As AICc is known to select more complex models (Sutherland et al., 2023), we also looked at how often the wrong model (here the proxy-informed model) would be selected if we only selected the proxy-informed model if ΔAICc *>* 2Δ*k*. Under this more conservative framework, the proxy-informed model was wrongly chosen only 2-4% of the time, which is close to the conventional 5% type I error rate.

### 3.3 Case studies

In case study 1, the RSF that included the interaction between the rate of mass gain of murres and distance to colony (AICc = 2980.33, *k* = 3) outperformed the baseline model (AICc = 2983.16, *k* = 2; ΔAICc = 2.82, Δ*k* = 1) and the coefficient associated with this interaction was significantly different from 0 (Table S2). This proxy-informed model indicated that, unlike individuals that lost mass, individuals that gained mass selected a range of foraging sites that included areas further from the colony (Figure 4B). In contrast to the baseline model that suggested that only areas very close to the colony were selected (Figure 4A), this proxy-informed model demonstrates that areas much further from the colony may be energetically beneficial (Figure 4D).

**Figure 4:**
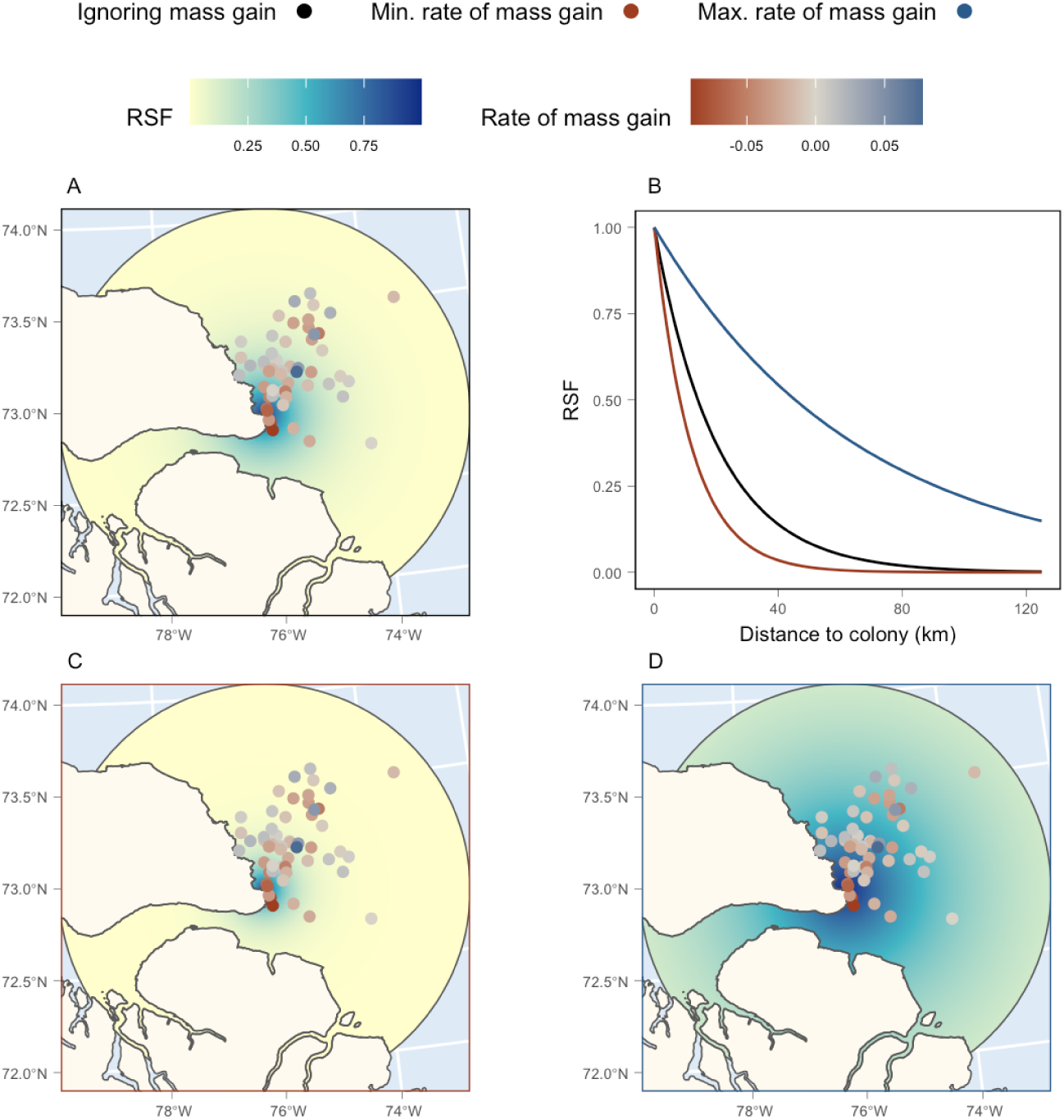
Results from the two murre RSF analyses. Panels A, C, and D show the predicted RSF value as a function of distance to colony for the baseline model, the proxy-informed model for the individual with the minimum rate of mass gained, and the proxy-informed model with the maximum rate of mass gained, respectively. The points represent the location for each foraging trip, with their colour representing the rate of mass gain. The colour scale of the circular study area represents the value of the predicted RSF, with darker colours representing higher selection. Panel B represents the predicted change in RSF as a function of distance to colony for the baseline model (black), the proxy-informed model for the individual with the minimum rate of mass gained (red), and the proxy-informed model for the individual with the maximum rate of mass gained (blue).

In case study 2, the SSF that included interactions between the number of gull chicks and both distance to colony and the land-human impact interaction (AICc = 522397.4, *k* = 13) outperformed the model that ignored this proxy of reproductive success (AICc = 522406.7, *k* = 11; ΔAICc = 9.35, Δ*k* = 2). Moreover, the coefficients associated with the number of chicks were significantly different from 0 (Table S4). According to this proxy-informed model, individuals with more chicks stayed closer to the colony and selected areas that were less impacted by humans, while individuals that had no chicks ranged further and selected for human-impacted landscapes (Figure 5). The baseline model characterised the habitat selection in a similar fashion to the proxy-informed model for individuals with one chick, with a moderate tendency to stay close to the colony and to select areas less impacted by humans.

**Figure 5:**
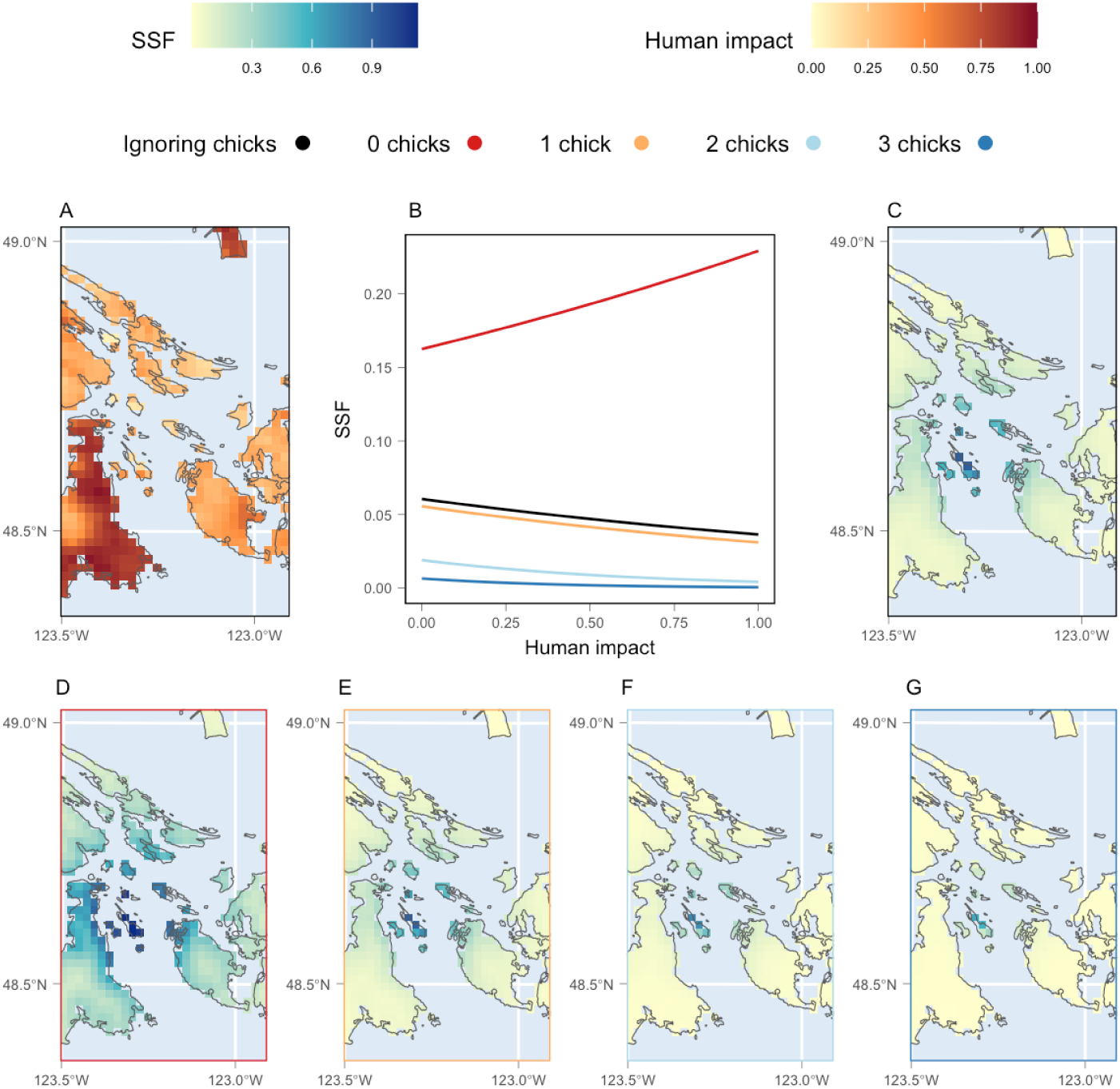
Results from the two gull SSF analyses. Panel A displays the cumulative human impact on land in the area close to the colony, with darker red colours representing more impacted landscapes. Panel B displays how the selection function changes in response to the cumulative human impact, with the black line representing the relationship from the model that ignores number of chicks, while the coloured lines represent the relationships from the model that includes interactions with the number of chicks (red = no chicks, orange = one chick, light blue = two chicks, and dark blue = three chicks). Panel C shows the spatial variation in the predicted SSF for the model that ignores the number of chicks, while panels D-G show the predicted SSF for the model that includes interactions with the number of chicks (D: no chicks, E: 1 chick, F: 2 chicks, G: 3 chicks). Dark blue colours represent higher SSF values.

In case study 3, the best SSF for narwhals included the presence of healed scars only in the movement kernel (AICc = 196139.0, *k* = 9), with a large difference in AICc from the baseline model (AICc = 196165.3, *k* = 8; ΔAICc = 26.27, Δ*k* = 1). The coefficient associated with the three-way interaction between healed scar presence, step length, and distance to shore was significantly different from 0 (Table S6). The next best model included the presence of healed scars in both the movement kernel and habitat selection (AICc = 196139.1, *k* = 10), but this more complex model had a relatively small difference in AICc from the best model (ΔAICc = 0.11, Δ*k* = 1) and the coefficient associated with the presence of healed scars on the selection for distance to shore was not significantly different from 0 (Table S6). Finally, the worst model was the one that included the presence of healed scars only in habitat selection (AICc = 196166.0, *k* = 9). Thus, the presence of healed scars appears to explain how narwhals alter their movement, but not their habitat selection. This lack of relationship with selection could be linked to the fact that distance to shore itself only appears to be an important covariate in the movement kernel (Table S6). Including the presence of healed scars in the movement kernel captured observed differences in mean step length; narwhals with healed scars moved slower far offshore than those without scars (Figure 6, Table S6). This difference in movement resulted in a distinction in areas used by narwhals, with individuals with healed scars spending more time further offshore than those without, despite little selection for these areas (Figures 6).

**Figure 6:**
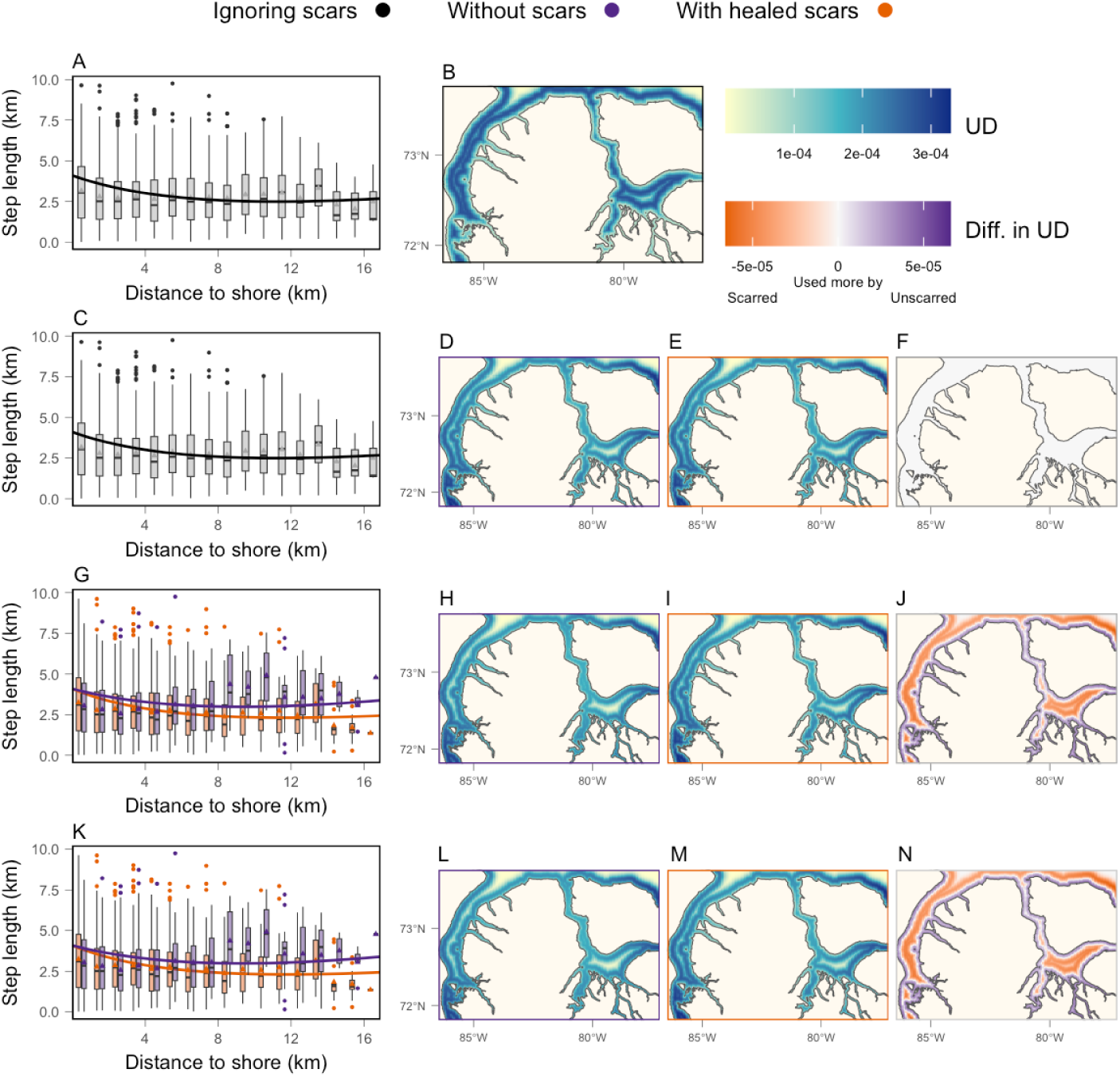
Results from the four narwhal SSF analyses: model that ignores scars (A-B), model that includes healed scars only in the habitat selection (C-F), model that includes healed scars only in the movement kernel (G-J), and model that includes healed scars in both the habitat selection and the movement kernel (K-N). Panels A, C, G, and K display the predicted mean step length, and boxplots showing the observed step lengths, with circles representing outliers and triangles representing the means. Panels B, D, E, H, I, L and M show the utilisation distribution (UD), with darker blue representing areas that are used more frequently. Panels F, J, and N display the difference between the UDs of unscarred (D, H, L) and scarred narwhals (E, I, M). In all panels, black represents model predictions or data when we ignore healed scars, while purple and orange represent model predictions and data for, or areas used more by, individuals without scars and with scars, respectively.

## 4 Discussion

Our literature review demonstrates that while habitat selection studies are often used to inform conservation actions, few recent studies link habitat selection to the types of fitness and health proxies that could help disentangle apparent habitat suitability from habitat quality. Our simulations emphasise how ignoring individual differences in health or fitness can bias habitat selection analyses. Specifically, they show how unbalanced samples, where more inferior individuals are tagged, can mislead researchers into inferring that good quality habitat is avoided, or that poor quality habitat is important. Our three case studies demonstrate the simplicity with which further insights into habitat selection can be gained by integrating health and fitness proxies in RSFs and SSFs.

The hidden relationships our case studies revealed by integrating individual performance covariates are ecologically meaningful and could impact conservation recommendations. Using the rate of mass gain, the RSF of case study 1 suggested that areas far from the colony, which would have been deemed secondary habitat by a naive RSF, may be energetically important for murres. These results likely reflect the central place foraging trade-off that has been observed in thick-billed murres: birds flying farther obtain larger prey (Elliott et al., 2009). The importance of such trade-offs for seabird conservation have been emphasised previously (Gaston et al., 2013). The SSF of case study 2 captured the central-place tendencies of breeding gulls and suggested that, although adults with low annual reproductive success select landscapes heavily impacted by humans, gulls successfully breeding in natural colonies avoid such areas. There is contradicting evidence regarding the value of urban landscapes and anthropogenic sources of food for chick growth and survival (e.g., Blight et al., 2015; Clewley et al., 2021; Kroc et al., 2024; Pais de Faria et al., 2021; van Donk et al., 2017). Although more detailed landscape layers are needed to identify the sources of their food (e.g., landfills vs mussel beds), our SSF appears to suggest that gulls in natural colonies select less-impacted areas during the early breeding period. Finally, case study 3 suggested that scarred narwhals move slower far from shore than unscarred individuals. Many studies relate narwhal behaviours to distance to shore (e.g., migration strategies, foraging-like movement; Dupont et al., 2025; Shuert et al., 2023), with some suggesting that narwhals respond to threats, such as killer whales and seismic surveys, by coming closer to shore (Breed et al., 2017; Heide-Jørgensen et al., 2021; Laidre et al., 2006). However, some sources of scars are more prominent close to shore (e.g., land-based hunting), and our results align with a previous analysis of this dataset that showed that narwhals with a higher composite stress index make smaller steps when displaying foraging-like behaviours far from shore (Shuert et al., 2025). Case study 3 demonstrates that movement itself, rather than selection, can change in relation to health and fitness proxies, and that individual differences in movement can affect the distribution of animals: scarred narwhals spent more time further from shore than unscarred individuals, despite not selecting for these areas. While our simple case studies demonstrate the ease with which a proxy can be included, and our results generally align with previous knowledge of these species, more comprehensive habitat selection analyses should be performed to draw any in-depth ecological conclusions or inform conservation planning.

The method we used to integrate health and fitness proxies is one of two approaches previously recommended for the inclusion of individual characteristics in habitat selection analyses (Northrup et al., 2022). The other is a two-step approach, in which a habitat selection analysis is applied to each individual and the estimated selection coefficients are subsequently compared (Bastille-Rousseau and Wittemyer, 2019; Winter et al., 2024). While both approaches have been used (Supporting Information) and each has advantages, neither can establish causation, nor whether changes in health/fitness lead to changes in habitat selection, or vice versa. We focused on using an interaction because of its simplicity; the only change in the analysis is adding one or more interactions with the proxy of interest. However, using a two-step approach can be equally valid. We are not promoting one approach over the other; our key message is that such proxies can be easily included in habitat selection analyses, and that doing so can provide crucial insights if interpreted carefully.

Many field programs measure useful health and fitness proxies, or are amenable to doing so in the future. Deploying tracking devices to collect data for RSFs and SSFs typically requires capturing animals, providing an opportunity to gather a range of useful information. At capture, many researchers: (1) perform health assessments, including screening for ectoparasites and injuries (e.g., Andrews et al., 2019; Gamblin et al., 2025; Lane et al., 2024; Shuert et al., 2025); (2) take biological samples that can be used to measure health-related variables such as stress hormone levels, contaminant loads, and oxygen storage capacity, (e.g., Aguilar et al., 2023; Andrews et al., 2019; Baak et al., 2024; Crossin et al., 2015; Shuert et al., 2025; Thompson et al., 2024); and, (3) measure mass and other useful morphometrics (e.g., Gamblin et al., 2025; Moreira-Arce et al., 2025; Thompson et al., 2024). Tracking devices that necessitate recapturing animals (e.g., GPS loggers; Nathan et al., 2022) offer unique opportunities to quantify how variables measured at capture have changed during the tagging period (e.g., change in mass; McComb-Turbitt et al., 2023; Patterson et al., 2025). Tracking systems that allow, or sometimes require, researchers to regularly resight tagged individuals (e.g., GPS logger combined with VHF or UHF) facilitate the monitoring of proxies such as reproductive success and survival (e.g., Alting et al., 2025; Lane et al., 2024; Parsons et al., 2025). Similarly, field programs that tag individuals at their breeding site facilitate short- to long-term monitoring of a variety of measures such as offspring count, food provisioning, and survival, using direct observation or automated systems (e.g., cameras, weighbridges; Beltran et al., 2023; Clewley et al., 2021; Elliott et al., 2009; Fijn et al., 2024; Lescroël et al., 2021). Studies using tracking devices concurrent with camera traps or drones could monitor individual performance metrics such as morphometrics, offspring counts, and dominance status (e.g., Alting et al., 2025; Kotik et al., 2023). Finally, movement data itself, or data collected via other sensors (e.g. acceleration, depth, heart rate) can provide information on energy gain and offspring survival (e.g., Adachi et al., 2023; Crossin et al., 2014; DeMars et al., 2013; Engebretsen et al., 2023; Patterson et al., 2025).

While collecting health and fitness data can increase costs, and the impacts of additional handling procedures should be considered carefully, their integration can provide the means to understand the inherent trade-offs of habitat selection. Many researchers have urged ecologists to link movement to fitness and health data (e.g., Gaillard et al., 2010; Hebblewhite and Haydon, 2010), with recent calls re-emphasising the need for auxiliary data such as trait values and demography (e.g., Beltran et al., 2025a; Beltran et al., 2025b). Along with these calls, there has been a growing push toward quantifying wildlife health (e.g., Aguilar et al., 2023; Aleuy et al., 2023; Kophamel et al., 2022), including the development of epigenetic metrics (Balard et al., 2024; Newediuk et al., 2025) and methods to combine multiple health metrics into a single overall measure of individual health, physiological dysregulation, or allostatic load (e.g., Edes et al., 2018; Milot et al., 2014; Shuert et al., 2025). Although long-term studies that quantify lifetime reproductive success (e.g., Beltran et al., 2023; McLoughlin et al., 2006) remain the gold standard for measuring fitness, many measures of individual health are linked to mortality risk (e.g., allostatic load; Edes et al., 2018), and frameworks that differentiate health determinants (e.g., infectious agents) from their processes (e.g., stress hormone production) and outcomes (e.g., survival; Aguilar et al., 2023) can help us distinguish habitats linked to these drivers of health.

We urge ecologists to collect and integrate fitness and health information in habitat selection analyses. Statistical methods to integrate them are available, and, in many cases, researchers simply need to use data they already collect. While each proxy must be interpreted carefully (e.g., annual reproductive success can be a poor metric for lifetime reproductive success), their use can provide insight into the causes and consequences of selection that would be otherwise difficult to gather. As such, we further encourage researchers to modify their field programs to systematically collect individual performance data. If fitness and health proxies are closely linked to overall fitness or lifetime reproductive success, including them in habitat selection analyses can bring us closer to identifying the habitats that promote reproduction and survival, and thus the habitats that should be the focus of conservation.

## 5 Acknowledgments

We thank the SȾÁUTW̱ (Tsawout) and W̱SIḴEM (Tseycum) for allowing access to X̱OX̱DEȽ (Mandarte Island) and Jade Baanstra, Amy Wilson, Peter Arcese, and Mark Hipfner for their support with fieldwork organisation and data collection. We thank the community of Mittimatalik (Pond Inlet) for its support in tagging operations and the devoted people who led operations in the field. We thank Jonathan Potts for help calculating the utilisation distribution. We thank the BC Knowledge Development fund, Canada Foundation for Innovation’s John R. Evans Leaders Fund, Canada Research Chairs program, Canadian Statistical Sciences Institute (CANSSI), Cecil and Kathleen Morrow Scholarship, Environment and Climate Change Canada (ECCC), Fisheries and Oceans Canada (DFO), Four Year Fellowship (4YF), Natural Sciences and Engineering Research Council of Canada (NSERC) Discovery Grant and Northern Supplement programs, Nunavut Implementation Fund, Nunavut Wildlife Management Board, Polar Continental Shelf Program, Werner and Hildegard Hesse Research Award in Ornithology award, Weston Family Awards in Northern Research, and World Wildlife Fund Canada for their financial support.

## Notes

https://github.com/MarieAugerMethe/InformedHSA

### Competing Interest Statement

The authors have declared no competing interest.

### Summary of Updates

Clarify some of the modelling terminology, update the literature review, and add available/control points

