## Supporting Information for "Including fitness and health proxies can alter our understanding of habitat selection"

### S1 Supporting Information

#### S1.1 Literature review

Our Web of Science search, conducted on April 28, 2025, returned 306 documents, of which 46 were duplicates and 14 were patents. From the 246 unique papers remaining, 90 were excluded based on the title, 45 based on the abstract, and 39 based on the full text. As such, the screening process left 72 papers to extract information from. From these 72 papers, we extracted the following information: (1) whether conservation or management inference was made based on the results; (2) whether it was an empirical or method-focused paper; (3) whether the habitat selection analysis included characteristics of individual fitness or health, and if so, which ones; (4) whether the habitat selection analysis included other individual characteristics (e.g., sex), and if so, which ones; and (5) if individual characteristics were included, the method used to do so. The information we extracted from the documents returned from the Web of Science search is available on GitHub: github.com/MarieAugerMethe/InformedHSA.

As mentioned in the main text, only three articles included a characteristic of individual health or fitness in their habitat selection analyses. These characteristics were: body size, reproductive status, and dominance status. One article included the characteristic as an interaction (as we have done in the main text), while the other two used a two-step approach.

#### S1.2 Simulations

##### S1.2.1 Resource selection function models

While the main text presents the models used to estimate the parameters of the resource selection functions (RSFs), here we present the general model for the RSFs themselves. We do not attempt to fully describe the theoretical framework that links these models; instead, we refer readers to references such as Matthiopoulos et al. (2023), Northrup et al. (2022), Warton and Shepherd (2010).

The RSFs used in the simulations and in the murre case study quantify the relative strength with which a unit is selected based on its covariates. We used the commonly adopted exponential form, which can be defined in general terms as:

$$w(\mathbf{x}) = \exp(\mathbf{x}^\mathsf{T}\boldsymbol{\beta}), \tag{S1}$$

where $\mathbf{x}^\mathsf{T}$ represents a row vector of covariates and $\boldsymbol{\beta}$ represents a column vector of selection coefficients. These functions are linked to the weighted distribution theory, which models the distribution of used habitats $f^u(\mathbf{x})$ as a function of the distribution of available habitat $f^a(\mathbf{x})$ (Northrup et al., 2022). Specifically, the distribution of used habitats is modelled as:

$$f^u(\mathbf{x}) = \frac{f^a(\mathbf{x})w(\mathbf{x})}{\int_{\mathbf{x}'\in\Omega} f^a(\mathbf{x}')w(\mathbf{x}')\mathrm{d}\mathbf{x}'}, \tag{S2}$$

where the denominator is an integral over the environmental domain $\Omega$ that ensures that $f^u(\mathbf{x})$ is a valid probability distribution function that integrates to 1. A common approach to estimate the parameters of an exponential RSF (equation S1) associated with this model (equation S2) is to use a logistic regression with data that includes a set of randomly generated available locations (Northrup et al., 2022). We adopted this widely used approximation in both the RSF simulations and the murre case study. As such, the equations presented in the main text (equations 2-3) and in the case study below (equations S6-S7) are the logistic regressions used to estimate the parameters, not the equations for the RSFs themselves. Thus, although the coefficients were estimated with a logit link function, predictions for the case study were made using the exponential form.

###### S1.2.2 Supplementary methods

While scenarios A and B are detailed in the main text, we expand here on the other simulation scenarios we explored. As briefly mentioned in the main text, we explore two step selection function (SSF) scenarios named scenarios C and D. These two scenarios were equivalent to scenarios A and B, respectively, except that we used SSFs rather than RSFs.

Step selection functions are similar to RSFs, but they account for the movement capacity of animals by limiting the distribution of available habitat to areas that can be reached between two successive locations (Avgar et al., 2016; Florko et al., 2025; Thurfjell et al., 2014). As such, the time-invariant distribution of available habitat found in the model associated with the RSF (equation S2) is replaced by a temporally varying distribution of available habitat. This time-varying availability distribution is often defined by a movement kernel. For a more detailed explanation of the relationship between RSFs and SSFs, please refer to Fieberg et al. (2021), Florko et al. (2025), and Northrup et al. (2022).

For each run of an SSF simulation, we simulated 100 steps for 30 individuals using the R package `amt` (Signer et al., 2019). The simulated movement model was based on the integrated SSF framework of Avgar et al. (2016), which models the movement as a combination of a habitat selection function and a selection-free movement kernel. The movement kernel describes the movement of the animal in the absence of resource selection. For all simulations, our selection-free movement kernel modelled the step length of the $n^{th}$ individual at time $t$ $l_{n,t}$ (i.e., distance between two consecutive locations) with a gamma distribution with shape and scale parameters equal to 1 and the turning angle of the $n^{th}$ individual at time $t$ $\theta_{n,t}$ (i.e., deviation from the previous bearing) with a von Mises distribution with kappa parameter equal to 1. We use the same selection parameters as for scenarios A and B: scenario C had $\beta_1 = 0.5$ and $\beta_2 = 1$, and scenario D had $\beta_1 = 0$ and $\beta_2 = -1$. We looked at the same three ratios of individuals and simulated each combination of scenario and proxy ratio 100 times for a total of 600 SSF simulations.

For each SSF simulation, we estimated the parameters of two SSFs: one baseline SSF that included only the environmental covariate, and one proxy-informed SSF with a selection function that included an interaction between the covariate and the health proxy. As for the RSF simulations, we adopted a widely used approximation approach to estimate the parameters of SSFs. As such, most of the equations presented below, as well as in the gull and narwhal case studies, are the models used to estimate the parameters, not the equations for the SSFs themselves. Specifically, to estimate the parameters of these two SSFs, we used the method described by Muff et al. (2020), which approximates the model likelihood by sampling control points and fitting a generalised linear mixed model (GLMM). For the baseline SSF, this method involved fitting a conditional Poisson regression:

$$\mu_{n,t,j} = \exp(\beta_{0,n,t} + \beta_{1,n} x_{n,t,j} + \\ \beta_l l_{n,t,j} + \beta_{\ln} \ln(l_{n,t,j}) + \beta_\theta \cos(\theta_{n,t,j})), \qquad y_{n,t,j} \sim \text{Poisson}(\mu_{n,t,j}), \tag{S3}$$

where $\beta_{0,n,t}$ is the step-specific intercept of animal $n$ at time $t$ that is modelled as a random effect with a fixed and large standard deviation value $\beta_{0,n,t} \sim \text{N}(0, 1000^2)$ and $\beta_{1,n}$ is the habitat selection coefficient and is modelled as a random effect $\beta_{1,n} \sim \text{N}(\beta_1, \sigma_1^2)$. The point $y_{n,t,j}$, covariate $x_{n,t,j}$, step length $l_{n,t,j}$, and turning angle $\theta_{ntj}$ are indexed by $j$ which represents whether it is a used step $y_{n,t,j} = 1$ for $j = 0$ or a control step $y_{n,t,j} = 0$ for $j = 1, \ldots, J$. Given that we have a predefined number of used and control points per step, the probability that the $n^{th}$ individual selects a point with covariate value $x_{n,t,j}$ given the set of possible covariate values $\mathbf{x}_{n,t} = (x_{n,t,0}, \ldots, x_{n,t,J})$ is:

$$\pi_{ntk} = \frac{\exp(\beta_1 x_{n,t,j} + \beta_l l_{n,t,j} + \beta_{\ln} \ln(l_{n,t,j}) + \beta_\theta \cos(\theta_{n,t,j}))}{\sum_{i=0}^{J} \exp(\beta_1 x_{n,t,i} + \beta_l l_{n,t,i} + \beta_{\ln} \ln(l_{n,t,i}) + \beta_\theta \cos(\theta_{n,t,i}))}. \tag{S4}$$

To estimate the parameters of the proxy-informed SSF model, we simply added an interaction with the health proxy:

$$\mu_{n,t,j} = \exp(\beta_{0,n,t} + \beta_{1,n} x_{n,t,j} + \beta_2 h_n x_{n,j,t} + \\ \beta_l l_{n,t,j} + \beta_{\ln} \ln(l_{n,t,j}) + \beta_\theta \cos(\theta_{n,t,j})), \quad y_{n,t,j} \sim \text{Poisson}(\mu_{n,t,j}), \tag{S5}$$

where $\beta_2$ is the coefficient that quantifies the interaction between health and selection for the environmental covariate. We sample 30 control points (J = 30) per used point using `amt` (Signer et al., 2019) for both models. We compared the two models using AICc.

As mentioned in the main text, we also performed two additional scenarios, where we simulated a baseline RSF (scenario E) and SSF (scenario F) model, with $\beta_1 = 0.5$, $\beta_2 = 0$. For scenario E, we followed the methods used for scenarios A and B. For scenario F, we followed the methods of scenarios C and D. In brief, we simulated the data for 30 individuals and did so 100 times. We fitted two models, the baseline model (simulated model) and the proxy-informed model with a non-relevant health proxy (wrong model). In these scenarios, the health proxy is not used to simulate the movement data. Thus, we could not explore different health value ratios.

###### S1.2.3 Supplementary results

In the main text, we detail the results of the RSF simulations (scenarios A and B); here, we present the equivalent results for the SSF simulations (scenarios C and D). As in scenario A, the estimates of the selection coefficient of the baseline SSF in scenario C ($\beta_1$ in equation S3) were negatively biased when there were more individuals in inferior health (Figure S1A), and as a result, their 95% confidence intervals overlapped with 0 in 93% of SSF simulations. Under a null hypothesis testing approach, these confidence intervals would be interpreted as evidence for a lack of selection of this covariate, even though individuals in good health select for it strongly. As in scenario B, the estimates of the selection coefficient of the baseline models ($\beta_1$ in equation S3) were positively biased when there were more unhealthy individuals in the sample (Figure S1B), and, as a result, their 95% confidence intervals excluded 0 in 55% of the SSF simulations. Under a null hypothesis testing approach, these confidence intervals would have been interpreted as evidence for selection for the environmental covariate, even though individuals in superior health condition avoid high values under these scenarios. Overall, as for the RSF simulations, when SSF simulations included a relationship with a health proxy, ignoring the health proxy biased the results when the ratio of individuals in different health conditions was unbalanced.

Figure S1: Boxplots of the estimated selection parameter $\beta_1$ of an SSF under two scenarios and three health ratios of individuals. The black horizontal lines represent the values used to simulate the data: $\beta_1 = 0.5$ and $\beta_1 = 0$ for scenario C (panel A) and D (panel B), respectively. Scenarios C and D have an increasing ($\beta_2 = 1$) and decreasing ($\beta_2 = -1$) selection for the covariate with increasing health/fitness proxy condition, respectively. The black boxplots represent the estimates when we used an SSF that ignored fitness proxies, while the pink boxplots represent the estimates when the fitness proxy was included in the SSF as an interaction with the environmental covariate.

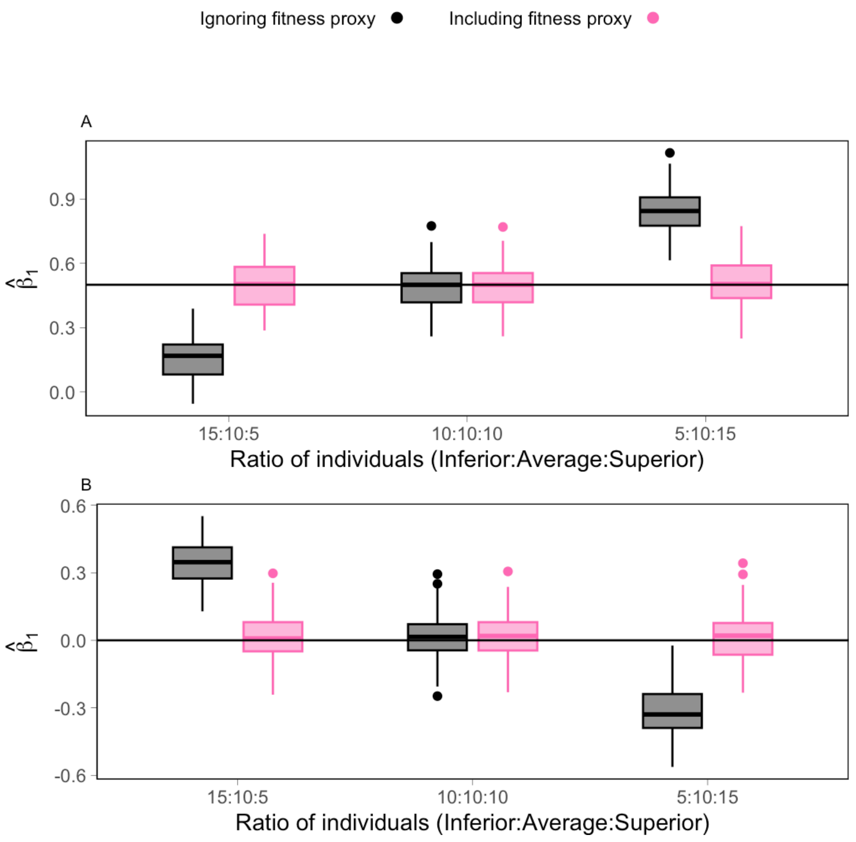


For both RSF and SSF simulations, the models that included the health proxy returned unbiased parameter estimates. The coverage was excellent for both $\beta_1$ and $\beta_2$, with the 95% confidence intervals of 95-96% of the RSF and SSF simulations containing the true simulated value. For the SSF simulations, the movement kernel did not appear to be affected by the inclusion of the health proxy in the habitat selection. All parameters of the movement kernel appeared unbiased, regardless of whether the health proxy was included in the model (Figure S2).

Figure S2: Boxplots of the estimated movement kernel parameters of SSFs under two scenarios and three health ratios of individuals. The black horizontal lines represent the value used to simulate the data: shape of gamma $\alpha_1 = 1$ (A-B), scale of gamma $\alpha_2 = 1$ (C-D), and kappa of von Mises $\kappa = 1$ (E-F). Panels A, C, and E display the results for scenario C with an increasing ($\beta_2 = 1$) selection for the covariate with increasing health proxy value. Panels B, D, and F display the results of scenario D with a decreasing ($\beta_2 = -1$) selection for the covariate with increasing health proxy value. The black boxplots represent the estimates when we used an SSF that ignored fitness proxies, while the pink boxplots represent the estimates when the fitness proxy was included in the SSF as an interaction with the environmental covariate.

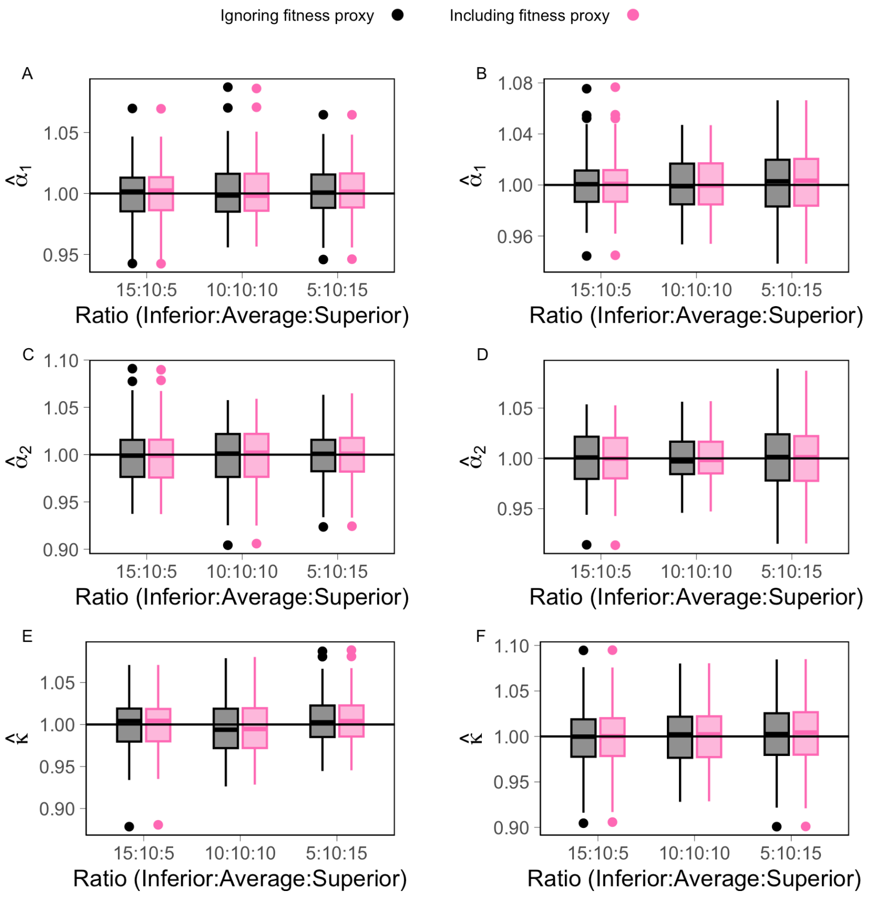

#### S1.3 Case study 1

##### S1.3.1 Supplementary methods

The movement data of thick-billed murres (hereafter murres) were collected in July and August 2022 and 2023 at the Cape Graham Moore colony on Bylot Island, Nunavut, Canada (72°56.5500 N, -76°5.6880 W). The murres were captured at the nest using a noose pole and weighed to the nearest gram. A Technosmart Axy-trek mini tag programmed to collect GPS locations at 1 or 3 min intervals was attached to the lower back feathers using Tesa tape, glue, and cable ties. We attempted to recapture all murres between 12 and 24 hrs later, so as to track birds over a single foraging bout. Our data pre-processing included: (1) restricting the data set to short deployments ($<$ 26 hrs) of incubating murres to limit tracks to single foraging bouts of adults that are not provisioning chicks; (2) choosing one foraging trip at random for each of the three individuals for which we had more than one to remove pseudo-replication; (3) removing two unrealistic locations that were more than 1000 km away from the colony; (4) excluding 16 locations that would have resulted from travelling at an unrealistic speed of 125 km/h or greater; and (5) standardizing the data set by subsampling the data collected every minute to keep only one location every three minutes. We then chose the location farthest from the colony to represent the destination of each foraging bout (i.e., where they likely forage before returning).

We created two RSF models for the data of 54 thick-billed murres (Table S1). The baseline model only included distance to colony as a covariate, while the proxy-informed model included an interaction with the rate of mass gained. To estimate the parameters of these

RSFs, we used the method of Muff et al. (2020), which approximates the model likelihood by sampling available locations and fitting a GLMM. The models described below are the GLMMs used to estimate the parameters of the RSFs, not the RSFs themselves.

##### Murre model 1 (baseline): rate of mass gain ignored

The model used to estimate the parameters of the baseline RSF is:

$$\text{logit}(\pi_{n,j}) = \beta_{0,n} + \beta_1 d_{n,j} \qquad y_{n,j} \sim \text{Bernoulli}(\pi_{n,j}), \tag{S6}$$

where logit represents the logistic link function, $y_{n,j}$ is a variable that represents whether point $j$ of individual $n$ is a used ($y_{n,j} = 1$ for $j = 0$) or available point ($y_{n,j} = 0$ for $j = 1, ..., J$), $d_{n,j}$ is the distance to colony for point $j$ of individual $n$, $\beta_1$ is the selection coefficient, and the intercept is modelled with a random effect that has a standard deviation fixed to a large value, here $\beta_{0,n} \sim \text{N}(\beta_0, 1000000^2)$.

##### Murre model 2 (proxy-informed model): rate of mass gain included

The model used to estimate the parameters of the proxy-informed RSF is:

$$\text{logit}(\pi_{n,j}) = \beta_{0,n} + \beta_1 d_{n,j} + \beta_2 m_n d_{n,j}, \qquad y_{n,j} \sim \text{Bernoulli}(\pi_{n,j}), \tag{S7}$$

which differs from murre model 1 (equation S6) only in one term $\beta_2 m_n d_{n,j}$, where $\beta_2$ is the interaction between selection for the covariate distance to colony $d_{n,j}$ and the rate of mass gain during the foraging trip $m_n$ for individual $n$.

We only have one location per trip and one trip per individual, so there is no need to

Table S1: Thick-billed murre data summary. For each individual (identified by their Canadian Wildlife Service bands), we present the date of capture, the duration of the deployment (in hours), the maximum distance from the colony for the farthest location of the foraging bout (in kilometres), the mass gained or lost during the deployment (in grams), and the rate of mass gain over the deployment (in grams per hour).

| Individual | Date of capture | Dist. from colony (km) | Duration (hr) | Mass gained (g) | Rate of mass gain (g/hr) |
|---|---|---|---|---|---|
| 96699530 | 2022-08-03 | 16.37 | 22.42 | -45 | -0.033 |
| 99698880 | 2023-07-25 | 33.53 | 15.23 | 4 | 0.004 |
| 99699507 | 2022-07-25 | 60.34 | 18.70 | -11 | -0.010 |
| 99699509 | 2022-07-25 | 70.89 | 16.42 | 28 | 0.028 |
| 99699510 | 2022-07-25 | 29.73 | 19.85 | -15 | -0.013 |
| 99699511 | 2023-08-01 | 32.66 | 15.68 | -16 | -0.017 |
| 99699513 | 2023-07-29 | 53.71 | 23.77 | -43 | -0.030 |
| 99699514 | 2023-08-01 | 23.80 | 14.05 | -20 | -0.024 |
| 99699516 | 2023-07-29 | 32.92 | 14.23 | 0 | 0.000 |
| 99699520 | 2023-08-01 | 32.21 | 10.32 | 48 | 0.078 |
| 99699521 | 2022-08-03 | 63.25 | 23.83 | -40 | -0.028 |
| 99699522 | 2023-08-01 | 6.36 | 11.35 | -54 | -0.079 |
| 99699523 | 2022-08-03 | 30.67 | 12.17 | 15 | 0.021 |
| 99699524 | 2022-08-03 | 67.30 | 22.35 | 1 | 0.001 |
| 99699525 | 2022-08-03 | 59.05 | 23.77 | -46 | -0.032 |
| 99699527 | 2022-08-03 | 28.14 | 12.27 | 11 | 0.015 |
| 99699528 | 2022-08-03 | 27.62 | 21.73 | -21 | -0.016 |
| 99699529 | 2022-08-03 | 4.40 | 10.88 | -23 | -0.035 |
| 99699533 | 2022-07-25 | 37.16 | 20.58 | -13 | -0.011 |
| 99699536 | 2022-07-25 | 72.65 | 16.07 | -7 | -0.007 |
| 99699544 | 2022-07-29 | 51.46 | 20.62 | -6 | -0.005 |
| 99699545 | 2022-07-29 | 43.13 | 14.75 | 10 | 0.011 |
| 99699546 | 2022-07-29 | 17.41 | 15.75 | 9 | 0.010 |
| 99699549 | 2023-08-02 | 47.72 | 13.37 | 8 | 0.010 |
| 99699550 | 2022-08-06 | 12.57 | 14.43 | -7 | -0.008 |
| 99699551 | 2022-08-06 | 16.73 | 20.58 | -24 | -0.019 |
| 99699553 | 2022-08-06 | 12.11 | 15.98 | 15 | 0.016 |
| 99699554 | 2022-08-06 | 15.27 | 12.77 | 1 | 0.001 |
| 99699556 | 2022-08-06 | 37.11 | 18.63 | 11 | 0.010 |
| 99699558 | 2022-08-06 | 31.77 | 11.33 | 11 | 0.016 |
| 99699559 | 2022-08-06 | 11.01 | 14.35 | -78 | -0.091 |
| 99699560 | 2023-08-01 | 10.80 | 15.48 | -45 | -0.048 |
| 99699561 | 2022-08-06 | 78.40 | 22.58 | 20 | 0.015 |
| 99699577 | 2022-08-08 | 2.59 | 22.88 | -93 | -0.068 |
| 99699578 | 2022-08-08 | 18.63 | 22.17 | -66 | -0.050 |
| 99699579 | 2022-08-08 | 45.76 | 22.92 | -15 | -0.011 |
| 99699580 | 2023-07-29 | 31.49 | 24.27 | -19 | -0.013 |
| 99699605 | 2023-08-02 | 46.22 | 23.20 | 8 | 0.006 |
| 99699654 | 2023-07-25 | 54.95 | 16.77 | 5 | 0.005 |
| 99699656 | 2023-07-25 | 105.91 | 25.43 | -24 | -0.016 |
| 99699661 | 2023-07-26 | 59.00 | 15.37 | -50 | -0.054 |
| 99699667 | 2023-07-29 | 38.10 | 23.98 | -46 | -0.032 |
| 99699671 | 2023-08-02 | 57.97 | 22.27 | -40 | -0.030 |
| 99699673 | 2023-08-01 | 27.96 | 11.13 | 5 | 0.007 |
| 99699674 | 2023-08-01 | 52.19 | 23.05 | -8 | -0.006 |
| 99699675 | 2023-08-01 | 34.63 | 12.32 | 30 | 0.041 |
| 99699677 | 2023-08-02 | 73.31 | 14.20 | 21 | 0.025 |
| 99699678 | 2023-08-02 | 57.05 | 13.67 | 36 | 0.044 |
| 99699679 | 2023-08-02 | 26.22 | 23.35 | -22 | -0.016 |
| 99699687 | 2023-07-29 | 3.41 | 12.88 | -44 | -0.057 |
| 99699688 | 2023-07-29 | 31.94 | 12.40 | -18 | -0.024 |
| 99699697 | 2023-08-03 | 26.10 | 25.52 | -47 | -0.031 |
| 130600360 | 2023-08-07 | 19.61 | 16.38 | -20 | -0.020 |
| 130600375 | 2023-08-06 | 48.80 | 23.45 | 18 | 0.013 |

account for pseudoreplication, and not enough data to quantify unexplained individual variation. Therefore, we have not included a random effect on $\beta_1$ (in murre models 1 and 2) nor $\beta_2$ (in murre model 2). However, we need a random effect on the intercept to account for the nested data structure. This nested data structure is created because we need to associate available points with each individual to assign them a rate of mass gain value so that we can include the rate of mass gain in the model. For each used point, we sampled 1000 available points from a study area defined by the water found in a circle of 125 km radius around the colony (Figure S3).

Figure S3: Used and available points used to estimate the parameters of the murre RSFs. The purple points are the location farthest from the colony for each foraging trip (i.e., used points). The grey points (largely overlapping with one another) are the available points that were sampled from the study area, which is defined as the water found within a 125 km radius of the colony.

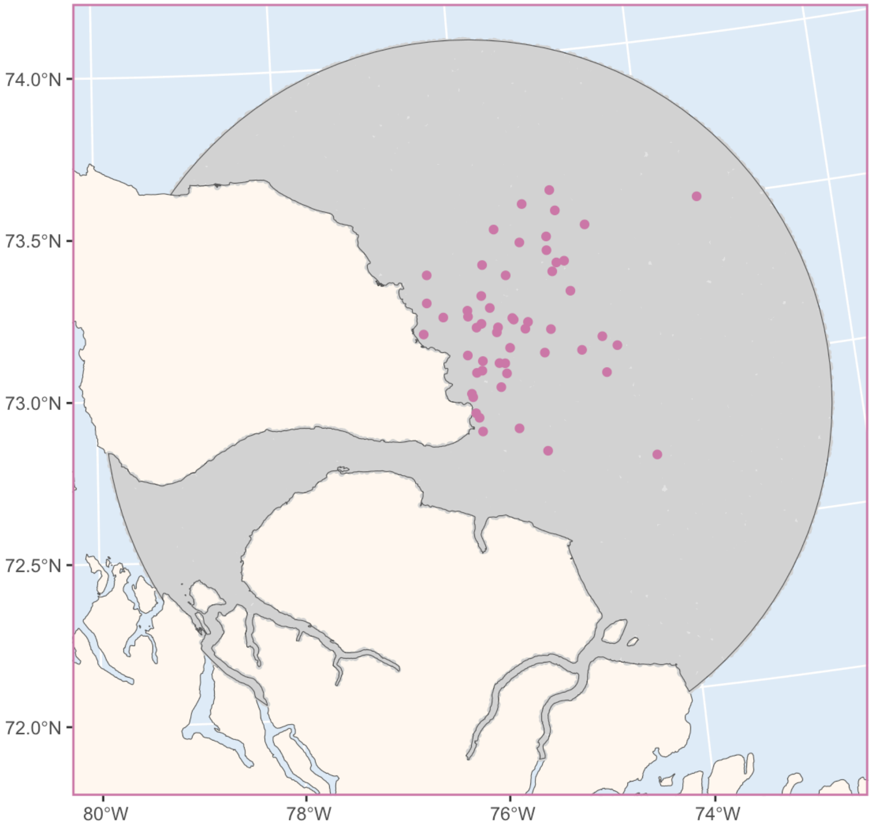

##### S1.3.2 Supplementary results

The parameter estimates from the murre RSFs are presented in Table S2. Note that we do not present the intercept value as it is not ecologically relevant, and is simply a function of the number of available per used points (Muff et al., 2020).

Table S2: Parameter estimates (with standard errors) for the two murre RSF models. A * indicates that the p-value comparing this parameter value to 0 is significant at the significance level of 0.05. Note that, as recommended by Muff et al. (2020), we have fixed the value of the standard deviation (SD) of the intercept to a large value (1e6). Note that while we estimate the intercept value, we do not report its value as it is ecologically irrelevant.

| Par: Variables | Rate of mass gain | |
|---|---|---|
| | Ignored | In selection function |
| $\beta_1$: $d_{n,j}$ | -0.04928* | -0.04691* |
| | (0.00548) | (0.00555) |
| $\beta_2$: $m_n \times d_{n,j}$ | | 0.408* |
| | | (0.191) |

#### S1.4 Case study 2

##### S1.4.1 Supplementary methods

Glaucous-winged gulls (hereafter gulls) were tagged on X̱OX̱DEȽ also known as Mandarte Island, in British Columbia, Canada (48°38.0400 N, -123°17.2200 W). The gulls were captured with noose mats that were constantly monitored. To limit impacts on captured gulls' reproductive success, we replace the eggs in the nest with wooden eggs during capture. We deployed OrniTrack-E20 4G tags (Ornitela) on 13 adult gulls captured at their nest during the early incubation period (June 11-13) of 2023. The Ornitela tag was attached using the leg-loop harness. Other measurements and samples were taken (e.g., blood, wing length), but were not used in this study. We removed one gull from the analysis because it was killed by an eagle eight days after release, resulting in 12 gulls. To monitor gulls and their nests, we placed a unique combination of Canadian Wildlife Service (CWS) and coloured plastic bands on their legs and a small numbered flag by their nest. We surveyed their nests every $6 \pm 4$ days (variation due to weather and field logistics) starting on June 14, 2023. Before the chicks began hatching, we walked to the nests to count eggs. After hatching, we primarily counted the chicks from blinds. We stopped monitoring at any sign of hatchling disturbance. The last date nests were monitored ranged from July 31 to August 2, and we used the number of chicks on those dates as our proxy of annual reproductive success. As hatchlings sometimes hid in nearby vegetation, including during the last chick survey, we combined information from previous surveys to identify the most likely number of chicks on July 31. If a tagged individual was not resighted at their nest, we assumed that it had forgone reproduction that year and assigned it a value of 0 chicks.

We used the cumulative human impact layer from Mu et al. (2021) and Mu et al. (2022). We used the layer for 2022, as this was the most recent layer available at the time of the analysis. Given that there is only one year difference between the movement data and this layer, and most pandemic restrictions were removed in early 2022, we trust this layer is a good approximation of the human impact in 2023. This raster layer provides data only for pixels that are on land, and land areas near the coast are sometimes given an NA value. As glaucous-winged gulls are coastal animals, we wanted to have information for all locations on land near the coast. As such, we interpolated values using the function `focal` of the R package `terra` (Hijmans, 2023). Specifically, we used the mean function with a weight equal to 9. Note that the human cumulative impact is only included as an interaction with the land binary covariate, so it is only used when the location is on land.

Table S3: Gull data summary. For each individual (identified by tag serial number), we present the date of capture, the number of days, the number of locations used in the analyses (only daytime locations outside a 200 m buffer from the colony), and the number of chicks they had on July 31, 2023.

| Individual | Date of capture | N. of days | N. of locations | N. of chicks |
|---|---|---|---|---|
| 223286 | 2023-06-12 | 40.4 | 1675 | 0 |
| 223287 | 2023-06-13 | 45.5 | 923 | 3 |
| 223290 | 2023-06-12 | 47.8 | 2265 | 0 |
| 223293 | 2023-06-11 | 48.8 | 1922 | 0 |
| 223295 | 2023-06-11 | 47.7 | 1121 | 0 |
| 223296 | 2023-06-11 | 47.3 | 929 | 1 |
| 223302 | 2023-06-12 | 47.6 | 1441 | 1 |
| 223305 | 2023-06-12 | 47.0 | 1190 | 1 |
| 223306 | 2023-06-13 | 46.6 | 1243 | 1 |
| 223307 | 2023-06-13 | 45.5 | 778 | 3 |
| 223315 | 2023-06-12 | 47.5 | 2853 | 1 |
| 223317 | 2023-06-12 | 47.6 | 849 | 0 |

We created two SSF models for the data of these 12 gulls (Table S3). These simple models were focused on how human impact on land may affect selection and accounted for the fact that these gulls are coastal animals that do not inhabit areas far inland or offshore and that they are central-place foragers during the incubation and chick-rearing period. The baseline model only included the environmental covariates, while the proxy-informed model included interactions with the number of chicks. To estimate the parameters of these SSFs, we used the method of Muff et al. (2020), which approximates the model likelihood by sampling control points and fitting a GLMM. The models described below are the GLMMs used to estimate the parameters of the SSFs, not the SSFs themselves.

#### Gull model 1 (baseline model): number of chicks ignored

The GLMM used to estimate the parameters of the baseline SSF models the point at the end of the step of individual $n$ at time $t$, $y_{n,t,j}$, which takes the value of 1 for used step ($j = 0$) and 0 for control points ($j = 1, \ldots, J$), as:

$$\begin{aligned} \mu_{n,t,j} = \exp(&\beta_{0,n,t} + \beta_{1,n} d_{n,t,j} + \beta_{2,n} c_{n,t,j} + \\ &\beta_{3,n} w_{n,t,j} + \beta_{4,n} w_{n,t,j} z_{n,t,j} + \\ &\beta_l l_{n,t,j} + \beta_{\ln} \ln(l_{n,t,j}) + \beta_\theta \cos(\theta_{n,t,j})), \qquad y_{n,t,j} \sim \text{Poisson}(\mu_{n,t,j}), \end{aligned} \tag{S8}$$

where for the $n^{th}$ individual at time $t$ for step $j$, $d_{n,t,j}$ is distance to shore, $c_{n,t,j}$ is the distance to colony, $w_{n,t,j}$ is whether the animal was on land (1) or not (0), $z_{n,t,j}$ is the cumulative human impact value. The coefficient $\beta_{0,n,t}$ is the step-specific intercept of animal $n$ at time $t$, which is modelled as a random effect $\beta_{0,n,t} \sim \text{N}(0, 1000000^2)$, while the coefficients $\beta_{1,n}$, $\beta_{2,n}$, $\beta_{3,n}$, and $\beta_{4,n}$ are the coefficients associated with the relationship with covariates and are modelled with random effects where we estimate the mean and standard deviation: $\beta_{1,n} \sim \mathrm{N}(\beta_1, \sigma_1^2)$, $\beta_{2,n} \sim \mathrm{N}(\beta_2, \sigma_2^2)$, $\beta_{3,n} \sim \mathrm{N}(\beta_3, \sigma_3^2)$, and $\beta_{4,n} \sim \mathrm{N}(\beta_4, \sigma_4^2)$. The rest of the equation is associated with the movement kernel, where $l_{n,t,j}$ and $\theta_{n,t,j}$ are the step length and turning angle of step $j$ of individual $n$ at time $t$, and $\beta_l$, $\beta_{\ln}$, and $\beta_\theta$ are coefficients to be estimated. Since the land variable, $w_{n,t,j}$ can only take two values: 0 and 1, it controls for the fact that human impact is only included when the animal is on land.

##### Gull model 2 (proxy-informed model): number of chicks included

The GLMM used to estimate the parameters of the proxy-informed SSF is based on the previous GLMM, but it includes interactions with the number of chicks.

$$
\begin{aligned}
\mu_{n,t,j} = \exp(&\beta_{0,n,t} + \beta_{1,n} d_{n,t,j} + \beta_{2,n} c_{n,t,j} + \\
&\beta_{3,n} w_{n,t,j} + \beta_{4,n} w_{n,t,j} z_{n,t,j} + \\
&\beta_5 r_n c_{n,t,j} + \beta_6 r_n w_{n,t,j} z_{n,t,j} + \\
&\beta_l l_{n,t,j} + \beta_{\ln} \ln(l_{n,t,j}) + \beta_\theta \cos(\theta_{n,t,j})), \qquad y_{n,t,j} \sim \mathrm{Poisson}(\mu_{n,t,j}),
\end{aligned} \tag{S9}
$$

which differ from gull model 1 only in two terms: $\beta_5 r_n c_{n,t,j}$ and $\beta_6 r_n w_{n,t,j} z_{n,t,j}$, which both include an interaction with the number of chicks $r_n$.

##### S1.4.2 Supplementary results

The parameter estimates for the gull SSFs are presented in Table S4.

Table S4: Parameter estimates (with standard errors) for the two gull SSFs. A * indicates that the p-value comparing this parameter value to 0 is significant at the significance level of 0.05.

| Par: Variables | Number of chicks | |
|---|---|---|
| | Ignored | In selection function |
| $\beta_1$: $d_{n,t,j}$ | -0.1726* | -0.1722* |
| | (0.0456) | (0.0458) |
| $\beta_2$: $c_{n,t,j}$ | -0.1065* | -0.0698* |
| | (0.0202) | (0.0220) |
| $\beta_3$: $w_{n,t,j}$ | 0.263 | 0.264 |
| | (0.214) | (0.217) |
| $\beta_4$: $w_{n,t,j}z_{n,t,j}$ | -0.509 | 0.344 |
| | (0.389) | (0.368) |
| $\beta_5$: $r_n c_{n,t,j}$ | | -0.0399* |
| | | (0.0162) |
| $\beta_6$: $r_n w_{n,t,j}z_{n,t,j}$ | | -0.927* |
| | | (0.268) |
| $\beta_l$: $l_{n,t,j}$ | -2.10e-05* | -2.11e-05* |
| | (4.28e-06) | (4.28e-06) |
| $\beta_{\text{ln}}$: $\ln(l_{n,t,j})$ | 0.000548 | 0.000573 |
| | (0.002959) | (0.002961) |
| $\beta_\theta$: $\cos(\theta_{n,t,j})$ | -0.0284* | -0.0284* |
| | (0.0121) | (0.0121) |
| $\sigma_1$: $\beta_1$ | 0.150 | 0.151 |
| $\sigma_2$: $\beta_2$ | 0.0673 | 0.0539 |
| $\sigma_3$: $\beta_3$ | 0.709 | 0.721 |
| $\sigma_4$: $\beta_4$ | 1.300 | 0.892 |

#### S1.5 Case study 3

##### S1.5.1 Supplementary methods

Case study 3 used narwhal movement data to demonstrate how habitat selection can be informed by external signs of trauma. We used data from a long-term monitoring program led by Fisheries and Oceans Canada (see Heide-Jørgensen et al., 2015; Orr et al., 2001; Shuert et al., 2021, for the handling protocol and tagging details); specifically, we used the data from the 14 individuals that were outfitted with TDR10 tags (Wildlife Computers, Inc.) that provided accurate Fastloc GPS data (Dujon et al., 2014). These narwhals were captured between July 31 and September 11, 2017 and August 17 and 18, 2018 in Tremblay Sound, Nunavut, Canada (72°21.1389 N, -81°05.855 W).

To account for the short-term effects of handling (Shuert et al., 2021), the migratory behaviour of narwhals (Shuert et al., 2022), and the irregular nature of Fastloc GPS for marine animals (Dujon et al., 2014), we pre-processed the movement data as in Shuert et al. (2025). First, we excluded locations within 24 hr and after 30 days of release, resulting in summer movement tracks of similar length. Second, as many SSF methods require data collected at regular time intervals (Signer et al., 2019), we used the R package `crawl` (Johnson and London, 2018; Johnson et al., 2008) to regularise the time series to a 1 hr interval. We split any track with more than 2 hrs and 5 min time gap into two and used crawl to predict track locations every 1 hr. We removed the 115 locations that crawl placed on land, and, as we need a minimum of three locations to calculate the turning angles used in the SSF, we removed tracks with fewer than three locations. These pre-processing procedures resulted in a total of 5886 locations (Table S5).

Table S5: Narwhal data summary. We present the date of capture, the number of days and locations used in the analyses, and whether the individual had healed scars. Individual 18001 had a shorter time series because its tag only transmitted for 10 days.

| Individual | Date of capture | N. of days | N. of locations | Healed scars |
|---:|---|---:|---:|---|
| 17001 | 2017-07-31 | 28.7 | 525 | FALSE |
| 17002 | 2017-07-31 | 28.5 | 529 | FALSE |
| 17003 | 2017-08-01 | 28.4 | 550 | TRUE |
| 17004 | 2017-08-03 | 29.0 | 491 | TRUE |
| 17005 | 2017-08-03 | 28.8 | 499 | TRUE |
| 17006 | 2017-08-03 | 26.5 | 462 | TRUE |
| 17007 | 2017-08-06 | 28.9 | 467 | TRUE |
| 17008 | 2017-08-13 | 28.4 | 538 | FALSE |
| 17011 | 2017-08-30 | 26.5 | 391 | TRUE |
| 17012 | 2017-09-02 | 27.8 | 443 | TRUE |
| 17013 | 2017-09-03 | 19.5 | 309 | FALSE |
| 17015 | 2017-09-04 | 27.1 | 197 | FALSE |
| 18001 | 2018-08-17 | 9.9 | 209 | TRUE |
| 18002 | 2018-08-17 | 28.6 | 276 | FALSE |

We created four SSFs for the narwhal data. The environmental covariate of interest for these SSFs was distance to shore, and we assumed that distance to shore at the end of a step affects the selection function, while distance to shore at the start of a step affects the movement kernel. To estimate the parameters of these SSFs, we used the method Muff et al. (2020), which approximates the model likelihood by sampling control points and fitting a GLMM. The models described below are the GLMMs used to estimate the parameters of the SSFs, not the SSFs themselves.

#### Narwhal model 1 (baseline model): scars ignored

The GLMM used to estimate the parameters of the baseline SSF models the observed locations of $n = 1, \ldots, 14$ narwhals at time $t$, encoded as $y_{n,t,j} = 1$ for $j = 0$, and the $J$

sampled control locations, encoded as $y_{n,t,j} = 0$ for $j = 1, \ldots, J$ as:

$$\begin{aligned}\mu_{n,t,j} = \exp(&\beta_{0,n,t} + \beta_{1,n} d_{n,t,j} + \\ &\beta_l l_{n,t,j} + \beta_2 d_{n,t-1,j} l_{n,t,j} + \\ &\beta_{\ln} \ln(l_{n,t,j}) + \beta_3 d_{n,t-1,j} \ln(l_{n,t,j}) + \beta_4 d^2_{n,t-1,j} \ln(l_{n,t,j}) + \\ &\beta_\theta \cos(\theta_{n,t,j})), \qquad y_{n,t,j} \sim \text{Poisson}(\mu_{n,t,j}),\end{aligned} \tag{S10}$$

where for the $n^{th}$ individual at time $t$ for step $j$, $d_{n,t,j}$ represents the distance to shore, $l_{n,t,j}$ the step length, and $\theta_{n,t,j}$ the turning angle. The coefficient $\beta_{0,n,t}$ is the step-specific intercept of animal $n$ at time $t$, which is modelled as a random effect $\beta_{0,n,t} \sim \text{N}(0, 1000000^2)$, while the coefficients $\beta_{1,n}$ is the coefficient associated with the habitat selection function and is modelled with a random effect for which we estimate the mean and standard deviation: $\beta_{1,n} \sim \text{N}(\beta_1, \sigma_1^2)$. The rest of the terms are associated with the movement kernel, where $\beta_2$, $\beta_l$, $\beta_{\ln}$, $\beta_\theta$, $\beta_3$, and $\beta_4$ are coefficients associated with the step lengths and turning angles. As explained in Avgar et al. (2016), the inclusion of the step length and its natural logarithm is a way to model the step length distribution as a Gamma distribution. Similarly, the inclusion of the cosine of the turning angle is a method to model the turning angle distribution as a Von Mises distribution. Here, $\beta_2$, $\beta_3$, and $\beta_4$ allow for the movement kernel, specifically the step length distribution, to vary as a function of distance to shore.

The three other models use healed scars as a health proxy by including interactions in the model with the variable $s_n$, which takes the value of 1 when healed scars are present on individual $n$ and 0 when individual $n$ has no healed scars.

#### Narwhal model 2: scar information in the selection function

The GLMM used to estimate the parameters of the SSF with scar information in the selection function uses as a base the narwhal model 1 described above. However, to allow for the selection function to differ between individuals with and without scars, we included an interaction between the presence of scars $s_n$ and the distance to shore at the end of the steps $d_{n,t,j}$:

$$\begin{aligned}\mu_{n,t,j} = \exp(&\beta_{0,n,t} + \beta_{1,n} d_{n,t,j} + \beta_5 s_n d_{n,t,j} + \\ &\beta_l l_{n,t,j} + \beta_2 d_{n,t-1,j} l_{n,t,j} + \\ &\beta_{\ln} \ln(l_{n,t,j}) + \beta_3 d_{n,t-1,j} \ln(l_{n,t,j}) + \beta_4 d^2_{n,t-1,j} \ln(l_{n,t,j}) + \\ &\beta_\theta \cos(\theta_{n,t,j})), \qquad y_{n,t,j} \sim \text{Poisson}(\mu_{n,t,j}),\end{aligned} \tag{S11}$$

where $\beta_5$ represents the coefficient quantifying the relationship between the presence of scars and distance to shore.

#### Narwhal model 3: scar information in the movement kernel

The GLMM used to estimate the parameters of the SSF with scar information in the movement kernel selection function uses as a base the narwhal model 1 described in equation S10. However, to allow for the relationship with speed and distance to shore to differ between individuals with and without scars, we include an interaction between the presence of scars $s_n$, step length $l_{n,t,j}$ and distance to shore at the start of the step $d_{n,t-1,j}$:

$$\begin{aligned}\mu_{n,t,j} = \exp(&\beta_{0,n,t} + \beta_{1,n} d_{n,t,j} + \\ &\beta_l l_{n,t,j} + \beta_2 d_{n,t-1,j} l_{n,t,j} + \beta_6 s_n d_{n,t-1,j} l_{n,t,j} + \\ &\beta_{\ln} \ln(l_{n,t,j}) + \beta_3 d_{n,t-1,j} \ln(l_{n,t,j}) + \beta_4 d^2_{n,t-1,j} \ln(l_{n,t,j}) + \\ &\beta_\theta \cos(\theta_{n,t,j})), \qquad y_{n,t,j} \sim \text{Poisson}(\mu_{n,t,j}),\end{aligned} \tag{S12}$$

where $\beta_6$ quantifies the relationship between the presence of scars, step length, and distance to shore.

##### Narwhal model 4: scar information in both the selection function and the movement kernel

The GLMM used to estimate the parameters of an SSF that allows the presence of scars to affect both the selection function and the movement kernel combines models 2 (equation S11) and 3 (equation S12):

$$\begin{aligned}\mu_{n,t,j} = \exp(&\beta_{0,n,t} + \beta_{1,n} d_{n,t,j} + \beta_5 s_n d_{n,t,j} + \\ &\beta_l l_{n,t,j} + \beta_2 d_{n,t-1,j} l_{n,t,j} + \beta_6 s_n d_{n,t-1,j} l_{n,t,j} + \\ &\beta_{\ln} \ln(l_{n,t,j}) + \beta_3 d_{n,t-1,j} \ln(l_{n,t,j}) + \beta_4 d^2_{n,t-1,j} \ln(l_{n,t,j}) + \\ &\beta_\theta \cos(\theta_{n,t,j})), \qquad y_{n,t,j} \sim \text{Poisson}(\mu_{n,t,j}).\end{aligned} \tag{S13}$$

##### S1.5.2 Supplementary results

The parameter estimates for the narwhal SSF models are presented in Table S6.

Table S6: Parameter estimates (with standard errors) for the four narwhal SSF models. A * indicates that the p-value comparing this parameter value to 0 is significant at the significance level of 0.05.

| Par: Variables | Healed scars | | | |
|---|---|---|---|---|
| | Ignored | In selection function | In movement kernel | In both |
| $\beta_1$: $d_{n,t,j}$ | -0.0876 | -0.0179 | -0.0836 | -0.000448 |
| | (0.0522) | (0.0768) | (0.0533) | (0.076873) |
| $\beta_l$: $l_{n,t,j}$ | 1.44e-04* | 1.44e-04* | 1.44e-04* | 1.44e-04* |
| | (1.95e-05) | (1.95e-05) | (1.94e-05) | (1.94e-05) |
| $\beta_2$: $d_{n,t-1,j} l_{n,t,j}$ | -4.21e-05* | -4.21e-05* | -2.84e-05* | -2.82e-05* |
| | (6.29e-06) | (6.29e-06) | (6.44e-06) | (6.44e-06) |
| $\beta_{\text{ln}}$: $\ln(l_{n,t,j})$ | 0.2589* | 0.2588* | 0.2503* | 0.2503* |
| | (0.0576) | (0.0576) | (0.0574) | (0.0574) |
| $\beta_3$: $d_{n,t-1,j} \ln(l_{n,t,j})$ | -0.0604* | -0.0602* | -0.0529* | -0.0528* |
| | (0.0239) | (0.0239) | (0.0239) | (0.0239) |
| $\beta_4$: $d^2_{n,t-1,j} \ln(l_{n,t,j})$ | 0.00735* | 0.00733* | 0.00738* | 0.00737* |
| | (0.00166) | (0.00166) | (0.00168) | (0.00168) |
| $\beta_\theta$: $\cos(\theta_{n,t,j})$ | 0.0213 | 0.0214 | 0.0221 | 0.0222 |
| | (0.0226) | (0.0226) | (0.0226) | (0.0226) |
| $\beta_5$: $s_n d_{n,t,j}$ | | -0.120 | | -0.143 |
| | | (0.101) | | (0.101) |
| $\beta_6$: $s_n d_{n,t-1,j} l_{n,t,j}$ | | | -2.42e-05* | -2.44e-05* |
| | | | (4.52e-06) | (4.52e-06) |
| $\sigma_1$: $\beta_{1,n}$ | 0.188 | 0.178 | 0.192 | 0.179 |
